# Recruitment of FBXO22 for Targeted Degradation of NSD2

**DOI:** 10.1101/2023.11.01.564830

**Authors:** David Y. Nie, John R. Tabor, Jianping Li, Maria Kutera, Jonathan St-Germain, Ronan P. Hanley, Esther Wolf, Ethan Paulakonis, Tristan M.G. Kenney, Shili Duan, Suman Shrestha, Dominic D.G. Owens, Ailing Pon, Magdalena Szewczyk, Anthony Joseph Lamberto, Michael Menes, Fengling Li, Dalia Barsyte-Lovejoy, Nicholas G. Brown, Anthony M. Barsotti, Andrew W. Stamford, Jon L. Collins, Derek J. Wilson, Brian Raught, Jonathan D. Licht, Lindsey I. James, Cheryl H. Arrowsmith

**Author notes:** These authors contributed equally: David Y. Nie, John R. Tabor. Co-corresponding authors: Lindsey I. James < >, Cheryl H. Arrowsmith < >. C4 Therapeutics, Watertown, MA, USA.

## Abstract

Targeted protein degradation (TPD) is an emerging therapeutic strategy that would benefit from new chemical entities with which to recruit a wider variety of ubiquitin E3 ligases to target proteins for proteasomal degradation. Here, we describe a TPD strategy involving the recruitment of FBXO22 to induce degradation of the histone methyltransferase and oncogene NSD2. UNC8732 facilitates FBXO22-mediated degradation of NSD2 in acute lymphoblastic leukemia cells harboring the NSD2 gain of function mutation p.E1099K, resulting in growth suppression, apoptosis, and reversal of drug resistance. The primary amine of UNC8732 is metabolized to an aldehyde species, which engages C326 of FBXO22 in a covalent and reversible manner to recruit the SCF^FBXO22^ Cullin complex. We further demonstrate that a previously reported alkyl amine-containing degrader targeting XIAP is similarly dependent on SCF^FBXO22^. Overall, we present a highly potent NSD2 degrader for the exploration of NSD2 disease phenotypes and a novel FBXO22-dependent TPD strategy.

## Introduction

Targeted protein degradation (TPD) is a proximity-based pharmacological strategy that harnesses the ubiquitin-proteasome system to degrade a protein of interest (POI).^1^ One common approach involves the use of heterobifunctional molecules, often referred to as proteolysis-targeting chimeras (PROTACs). PROTACs consist of two binding moieties, one of which engages an E3 ubiquitin ligase and the other a POI. The two ligands are tethered to one another by a linker that can vary in structure and length. An effective chemical degrader recruits an E3 ligase into close proximity with the target POI to form a ternary complex, resulting in polyubiquitination of the POI and proteasome-mediated protein degradation.

There has been an explosion of interest in TPD technologies in the past two decades, enabling the degradation of an increasing number of protein targets.^1^ However, despite this growth, the parallel expansion of recruitable E3 ubiquitin ligases has not occurred at the same rate. In fact, the vast majority of targeted protein degraders act through the recruitment of one of two E3 ligases, VHL or cereblon (CRBN), which is striking considering the human genome encodes over 600 different E3 ligases.^2–4^ This limitation presents a number of obstacles for the TPD field. First, some cell types lack expression of VHL and CRBN, and thus, these E3 ligases are not universally recruitable in all disease states.^3,4^ For example, VHL is downregulated in certain malignancies, which limits the use of TPD as a therapeutic solution in such contexts.^5,6^ Moreover, many protein targets have remained recalcitrant to the traditional plug-and-play strategy of simply appending a known CRBN or VHL ligand to a selective ligand for a POI through a range of linker moieties. This may be due to the failure to form a *productive* ternary complex to enable the transfer of ubiquitin to an accessible lysine residue on the target protein to promote degradation. In addition to these general limitations, each E3 ligase can pose its own unique challenges. For example, VHL recruiting ligands are antagonists of native VHL protein interactions; therefore, hijacking VHL for TPD applications may be undesirable, given that VHL is a well-known tumor suppressor. CRBN-recruiting ligands, such as lenalidomide, thalidomide, and pomalidomide, are known immunomodulatory agents that have neo-substrates, including SALL4, IKZF1, and IKZF3^7^, which could result in off-target degradation. Finally, many TPD hetero-bifunctional compounds are relatively large molecules, exceeding the size of traditional small-molecule drugs. This can lead to poor cell permeability and pharmacological properties, necessitate extensive optimization, and limit their use clinically. To overcome these challenges, the availability of a diverse chemical toolkit of E3 recruiters will be critical for TPD to reach its full potential.

We recently reported UNC8153, an effective chemical degrader of NSD2 (nuclear receptor-binding SET domain-containing 2), which is a histone methyltransferase that dimethylates lysine 36 of histone 3 (H3K36me2), an epigenetic mark associated with active gene transcription.^8,9^ Mutations at the *NSD2* locus^8,10,11^ lead to overexpression and/or gain-of-function activity of NSD2, which leads to the upregulation of H3K36me2 in multiple myeloma and acute lymphoblastic leukemia, among other cancers, and are correlated with disease progression.^12^ UNC8153 potently degrades NSD2 in cells with exquisite selectivity and results in a significant reduction of the H3K36me2 mark in NSD2 mutant cells while also influencing certain disease-associated phenotypes in multiple myeloma cells.^9^ UNC8153 makes use of a simple primary alkylamine as its E3 recruiting moiety, resulting in a lower molecular weight relative to traditional heterobifunctional degraders. Interestingly, UNC8153-mediated degradation of NSD2 is both proteasome- and neddylation-dependent, hinting that the putative E3 ligase recruited by UNC8153 may be a component of one or more of the Cullin-RING E3 ligase complexes. Here, we report the discovery of UNC8732, a second-generation NSD2 degrader that demonstrates improved NSD2 degradation and H3K36me2 reduction in cells. Similar to UNC8153, UNC8732 also contains a primary alkyl amine, which we demonstrate is metabolized in cells to an active aldehyde species that promotes NSD2 degradation through the recruitment of FBXO22, a substrate recognition subunit of a Cullin 1 E3 ligase complex. Finally, we demonstrate that a previously reported primary amine-containing chemical degrader targeting the E3 ligase XIAP^13^ also acts through an FBXO22-dependent mechanism. Excitingly, this study expands the repertoire of strategies available for the development of new TPD reagents and therapeutics while also providing a valuable chemical probe for the modulation of NSD2 and H3K36me2.

## Results

### Discovery of UNC8732

With the goal of improving the efficacy of our first-generation NSD2 degrader, UNC8153 (DC_50_ = 0.35 µM), we focused on modifying the linker region between the NSD2 binding unit^14^ and the primary amine. We observed that shortening and lengthening the linker by a single methylene group, as in UNC8269 and UNC8325, respectively, resulted in somewhat weaker DC_50_ values (concentration at which 50% of NSD2 is degraded) as determined by an in-cell western (ICW) assay, suggesting that an eight-atom linker between the amine and the rightmost phenyl ring of the NSD2 ligand is optimal (Table 1).^9^

**Table 1:**
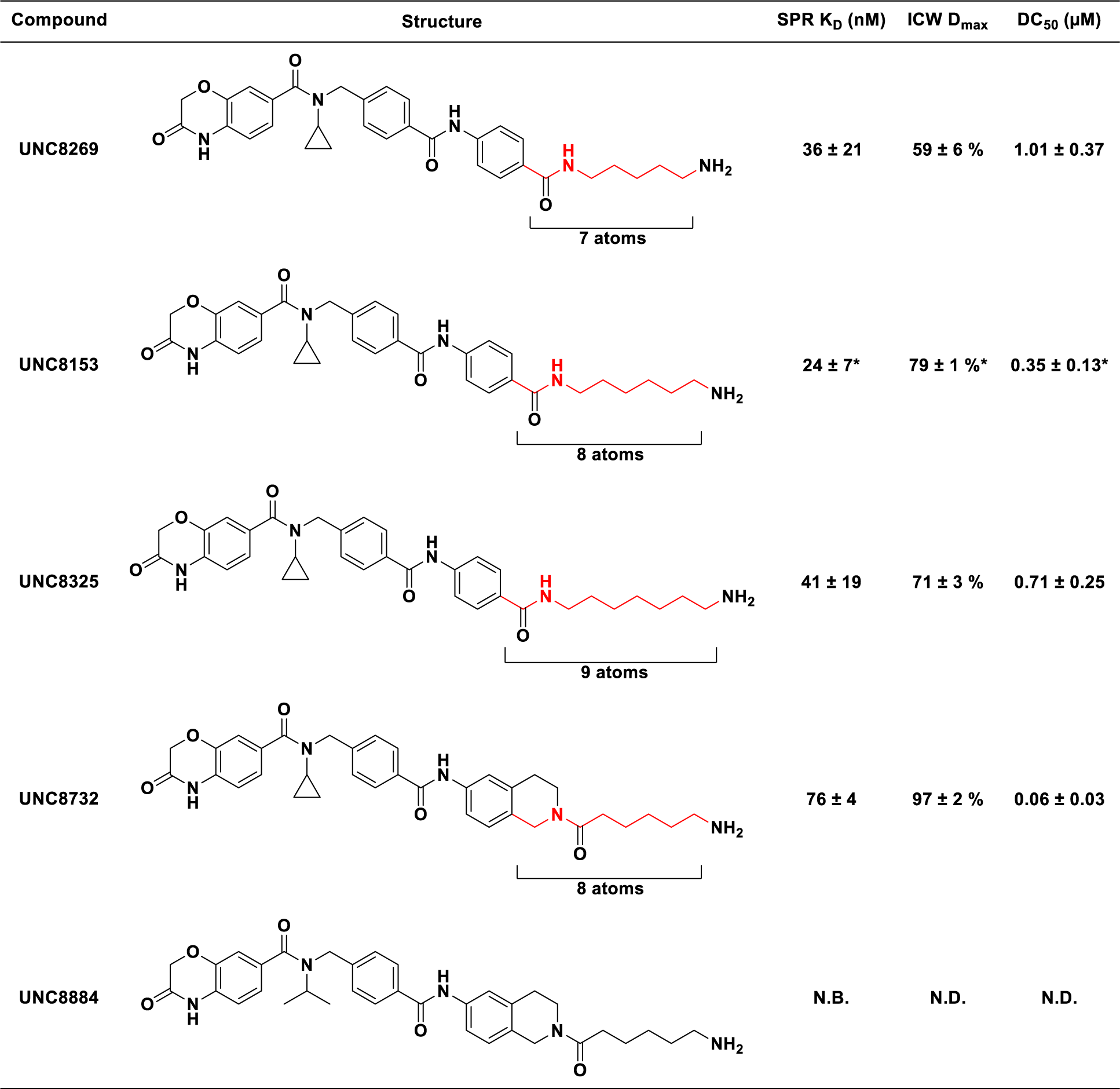
Discovery of UNC8732. Structures of NSD2 degraders with varying linker lengths. Their associated K_d_ values for NSD2-PWWP1 determined by SPR, D_max_, and DC_50_ values determined by an ICW assay are reported. Values reported as the average of 3 independent experiments ± standard deviation (SD). * denotes previously reported values in Hanley et al.^9^ N.B. denotes no binding up to 2 µM. N.D. denotes not determined.

| Compound | Structure | SPR $K_D$ (nM) | ICW $D_{max}$ | $DC_{50}$ (μM) |
| --- | --- | --- | --- | --- |
| UNC8269 | | $36 \pm 21$ | $59 \pm 6 \%$ | $1.01 \pm 0.37$ |
| UNC8153 | | $24 \pm 7^*$ | $79 \pm 1 \%^*$ | $0.35 \pm 0.13^*$ |
| UNC8325 | | $41 \pm 19$ | $71 \pm 3 \%$ | $0.71 \pm 0.25$ |
| UNC8732 | | $76 \pm 4$ | $97 \pm 2 \%$ | $0.06 \pm 0.03$ |
| UNC8884 |  | N.B. | N.D. | N.D. |

Due to the flexible nature of the linker, we next explored if we could “lock” the linker into a slightly more rigid and favorable conformation by constraining the amide linkage. Using this strategy, we discovered UNC8732, which contains a fused bicyclic ring system and is a highly potent NSD2 degrader with a DC_50_ of 0.06 ± 0.03 µM and a D_max_ of 97 ± 2% (Table 1). As UNC8732 does not bind NSD2-PWWP1 more potently than UNC8153, this improvement in degradation efficiency is not the consequence of improved binding to NSD2. For cellular studies, we also synthesized a negative control compound, UNC8884, which significantly reduces NSD2 binding and, therefore, does not promote NSD2 degradation (Figure 1A).

**Figure 1.**
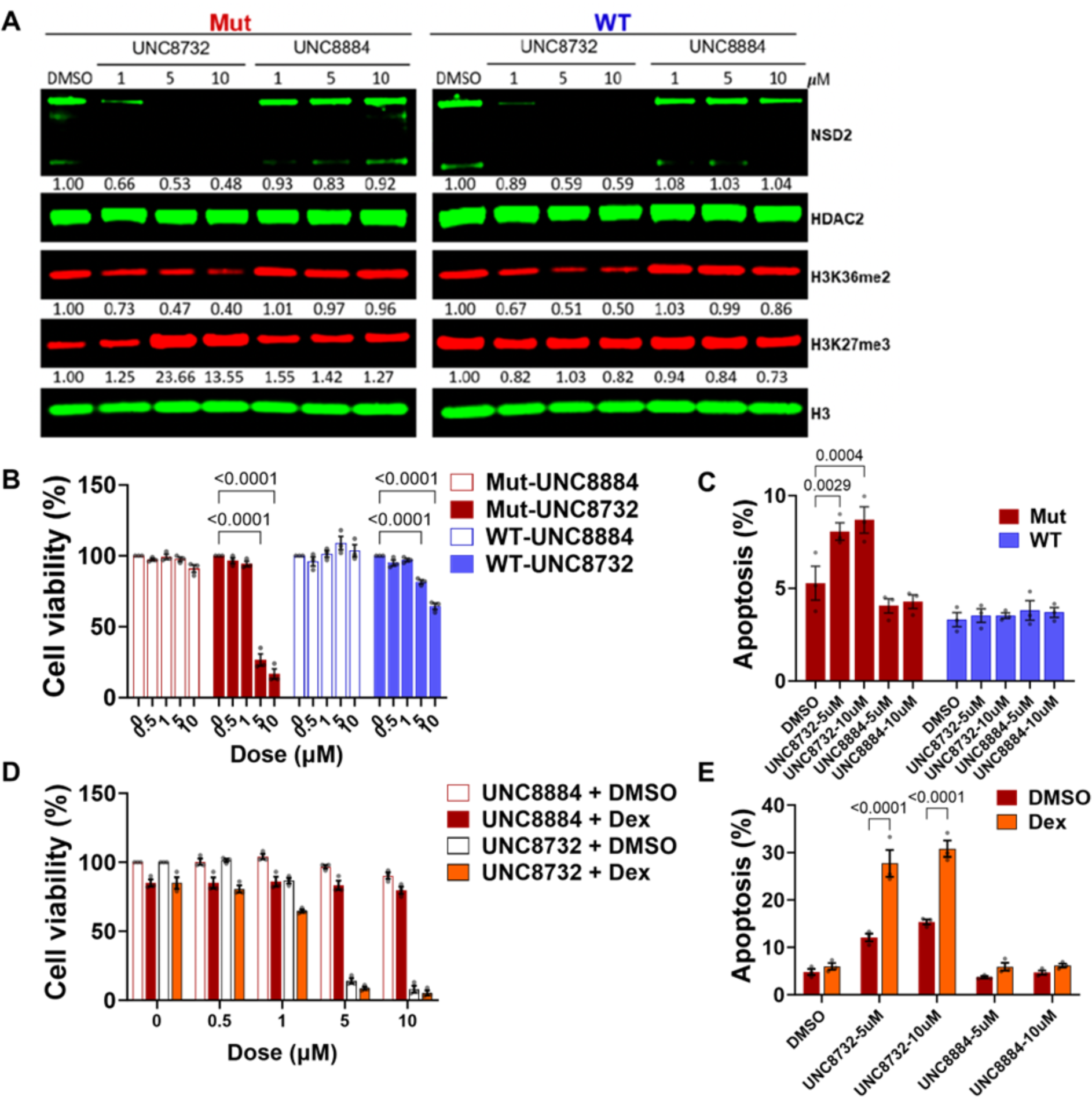
UNC8732 inhibits cell growth and restores glucocorticoid sensitivity of *NSD2* p.E1099K mutant ALL cells. **(A)** RCH-ACV cells were treated for 11 days with 0-10 µM UNC8732 or UNC8884. Cells were collected and analyzed by immunoblotting. H3 and HDAC2 are loading controls. NSD2 protein level is normalized to HDAC2. H3K36me2 and H3K27me3 levels are normalized to H3. **(B)** Viability of isogenic RCH-ACV ALL cell lines determined by CellTiter-Glo after treatment with varying concentrations of UNC8732 and UNC8884 for 18 days. **(C)** Apoptosis of isogenic RCH-ACV cell lines detected using Annexin V/PI staining by flow cytometry after treatment with varying concentrations of UNC8732 and UNC8884 for 18 days. **(D)** Viability of *NSD2* mutant RCH-ACV cells determined by CellTiter-Glo after pretreatment with varying concentrations of UNC8732 and UNC8884 for 18 days, followed by dexamethasone (1 µM) for 72 hours. **(E)** Apoptosis of *NSD2* mutant RCH-ACV cells detected using Annexin V/PI staining by flow cytometry after pretreatment with varying concentrations of UNC8732 and UNC8884 for 18 days followed by dexamethasone (1 µM) for 72 hours. Error bars represent mean ± SEM from three biological replicates. The statistical significance was evaluated using the Two-way ANOVA test and the p-values are displayed. WT, NSD2 WT; Mut, *NSD2* p.E1099K; Dex, Dexamethasone.

### UNC8732 triggers cell death and reverses glucocorticoid resistance in NSD2 mutant ALL

We previously demonstrated that the *NSD2*p.E1099K mutation drives oncogenic reprogramming^15^ and glucocorticoid (GC) resistance, potentially contributing to relapse of pediatric acute lymphoblastic leukemia (ALL).^16^ The p.E1099K mutation, due to its increased enzymatic activity^17,18^, leads to an increase of H3K36me2 chromatin mark across the genome of ALL cells and alteration of transcriptional programs.^19^ Since our NSD2 degraders effectively reduce global cellular levels of H3K36me2,^9^ we hypothesized that UNC8732 might reverse the aberrant H3K36me2 levels and associated oncogenic transcriptional program in ALL cells with the *NSD2* p.E1099K mutation. Here, we treated isogenic *NSD2* p.E1099K and *NSD2* wild type (WT) RCH-ACV cell lines^19^ with UNC8732 or negative control UNC8884 for up to 21 days, over which the chromatin marks slowly change. At 11 days, UNC8732 had dramatically degraded NSD2 protein in both *NSD2* mutant and WT RCH-ACV cell lines with concomitant reduction in the level of H3K36me2 in both *NSD2* mutant and WT RCH-ACV cell lines by >50% (**Figure 1A**). In *NSD2* p.E1099K mutant cells, which display high levels of H3K36me2 and low levels of the transcriptionally repressive H3K27me3 mark relative to *NSD2* WT cells, this epigenetic pattern was reversed by NSD2 degradation, with cells displaying both decreased H3K36me2 and a striking increase in H3K27me3 (**Figure 1A**). No change in histone marks occurred with UNC8884 treatment. Over the course of treatment (cells monitored at 12, 15, 18, and 21 days), the ALL cell lines displayed a dose-dependent and time-dependent decrease in viability in response to UNC8732 but not UNC8884 (**Figures 1B, Extended Figure 1A-C)**. This effect was much more pronounced in *NSD2* p.E1099K cells, which exhibited a greater than 50% reduction in cell viability at doses of 5 µM and 10 µM. With treatment of UNC8732 (10 µM), the viability of *NSD2* mutant cells decreased to 16.7% at day 18 compared with 64% for WT cells (Figure 1B), and at 21 days, viability decreased to 8% for *NSD2* mutant cells but only to 50% for *NSD2* WT cells (**Extended Data Figure 1C)**. At all-time points, UNC8732 induced apoptosis of *NSD2* mutant cells, with up to 15% positive for Annexin V at day 21, while significant apoptosis was not observed in degrader-treated *NSD2* WT cells (**Figure 1C, Extended Figure 1D-H**).

*NSD2* mutation and hyperactivity deregulates the epigenome of ALL, leading to aberrant repression of the *NR3C1* gene encoding the glucocorticoid receptor.^19^ We, therefore, determined whether NSD2 degradation in *NSD2* p.E1099K mutant ALL cells could restore GC sensitivity. *NSD2* mutant and WT ALL cells were pre-treated with UNC8732 or UNC8884 for 18 days, followed by combination treatment with dexamethasone (a GC drug) for 3 days. Dexamethasone itself induced no more than a 20% reduction in cell viability in *NSD2* mutant RCH-ACV cells and failed to induce apoptosis, while in the presence of the UNC8732, dexamethasone added to the effect of NSD2 degradation to reduce cell viability and induced apoptosis of *NSD2* mutant cells (**Figure 1D-E, Extended Data Figure 1I-O**). After 18 days, ∼15% of cells were induced into apoptosis by UNC8732 (10 µM) alone, while combination treatment led to >30% cell death (**Figure 1E, Extended Data O**). These data suggest that degradation of NSD2 by UNC8732 reverses the aberrant epigenetic landscape in *NSD2* mutant ALL cells, relieving GC resistance and potentially other aggressive biological properties resulting from the *NSD2* p.E1099K mutation.

### Metabolism of UNC8732 to the corresponding aldehyde drives NSD2 degradation

With promising results using our NSD2 degrader in a therapeutically relevant disease system but a limited understanding of how UNC8732 and related analogs promote NSD2 degradation, we sought to confirm the identity of the active compound in cells. Given that a primary alkylamine can be a metabolically labile group in a cellular context, it remained a possibility that the amine could be further metabolized into a different active species. We previously synthesized and evaluated several potential metabolites of the primary amine of UNC8153, none of which were able to effectively degrade NSD2, thereby excluding them from involvement in the mechanism of degradation^8^; however, not all possible metabolites were accessible synthetically. We conducted a liquid chromatography-mass spectrometry (LCMS)-based metabolite identification study with our first-generation degrader, UNC8153. Briefly, U2OS cells were treated with 10 µM of UNC8153 for 24 hours, and the components of the resulting lysates were analyzed by LC-MS/MS. Not surprisingly, a metabolite was identified that corresponded to the conversion of the primary amine to an aldehyde (Figure 2A top). To further validate this observation, U2OS cells in DMEM culturing media and 10% Fetal Bovine Serum (FBS) were treated with our second-generation degrader, UNC8732 (10 µM), and the resulting lysate was analyzed by LC-MS/MS. Similarly, the mass of the aldehyde derivative of UNC8732 was observed (Figure 2A bottom, 2B). The concentration of UNC8732 rapidly decreased, dropping below the limit of detection within 6 hours, while the aldehyde metabolite formed in significant quantities after 6 hours.

**Figure 2:**
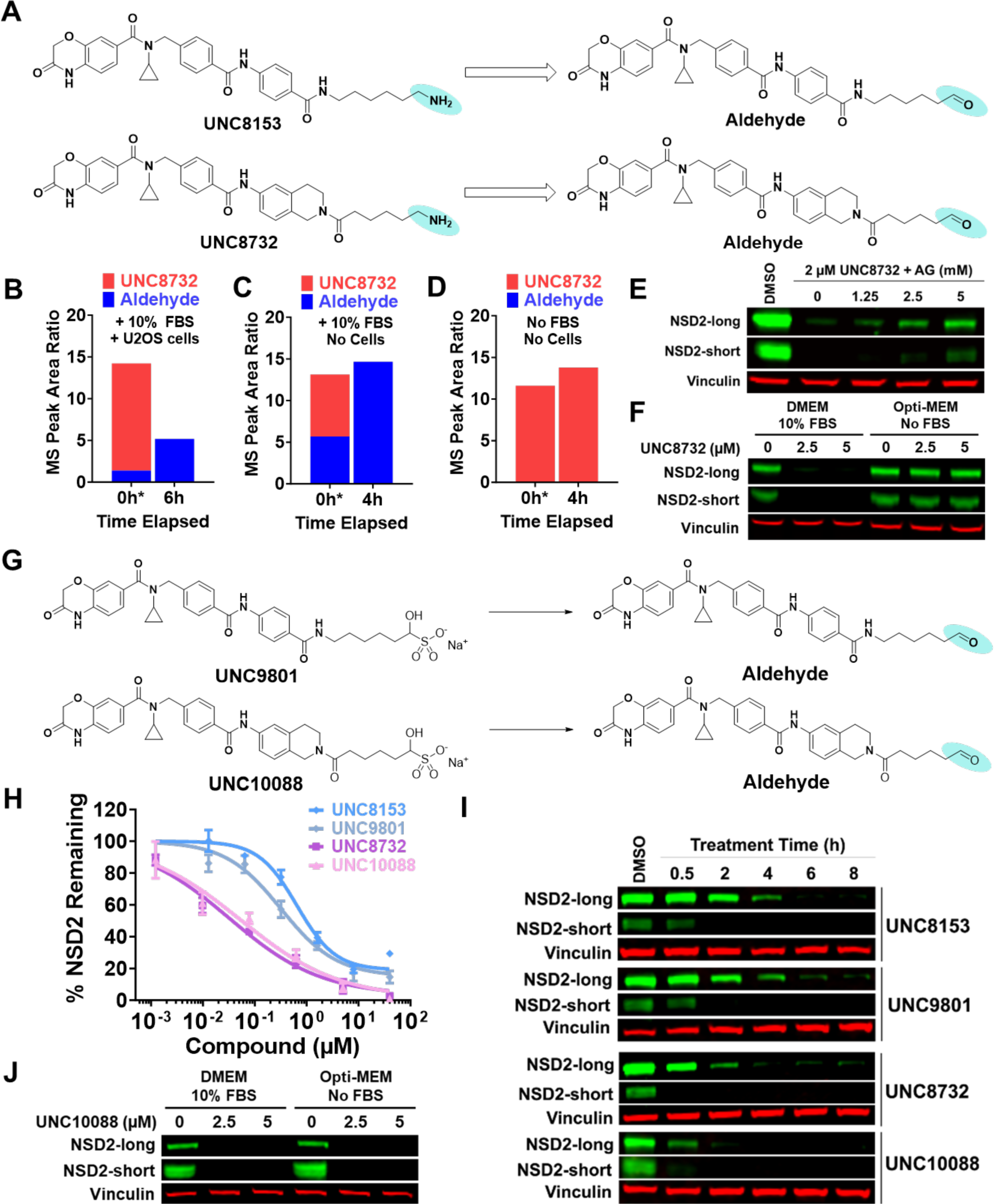
UNC8732 metabolism to the corresponding aldehyde drives NSD2 degradation. (A): Compound structures of UNC8153 (top) and UNC8732 (bottom) prodrugs and their corresponding aldehyde metabolites. (B): MS peak area ratio representing the relative levels of UNC8732 and its aldehyde metabolite in lysates generated from U2OS cells treated with UNC8732 and cultured with DMEM and 10% FBS at indicated time points. *For Figure 2B-D, 0 hours denotes mass spectrometric analysis performed immediately after compound treatment, which includes a very brief sample preparation time to process for MS. (C): MS peak area ratio representing the relative levels of UNC8732 and its associated aldehyde species in cell-free DMEM and 10% FBS at indicated time points. (D): MS peak area ratio representing the relative levels of UNC8732 and its associated aldehyde species in cell-free DMEM without FBS at indicated time points. (E): U2OS cells were treated with 2 µM UNC8732 and varying concentrations of aminoguanidine (AG) for 6h. NSD2 levels were measured by immunoblot. The experiment was performed 3 times with consistent results. Vinculin is a loading control. (F): U2OS cells were cultured in DMEM + 10% FBS or Opti-MEM with no FBS while being treated with the indicated concentrations of UNC8732 for 24h, followed by immunoblotting analysis for NSD2 levels. The experiment was performed 3 times with consistent results. Vinculin is a loading control. (G): Compound structures of UNC9801 (top) and UNC10088 (bottom) prodrugs and their corresponding hydrolyzed aldehyde products. (H): U2OS cells were treated with the specified compounds in a dose-response format for 24h. NSD2 levels were measured by an ICW assay. The data presented are from one representative experiment, including 4 technical replicates. (I): U2OS cells were treated for the indicated amount of time with UNC8732 (2 µM), UNC10088 (2 µM), UNC8153 (10 µM), or UNC9801 (10 µM), and were subsequently analyzed by immunoblotting. Vinculin is a loading control. The experiment was performed 3 times with consistent results. The data shown are representative immunoblots. (J): Cells were cultured in DMEM + 10% FBS or Opti-MEM with no FBS and were subsequently treated with indicated concentrations of UNC10088 for 24h, followed by immunoblotting analysis for NSD2 levels. The experiment was performed 3 times with consistent results. Vinculin is a loading control.

We next incubated DMEM media containing FBS with UNC8732 in the absence of cells and again observed the formation of the corresponding aldehyde metabolite, while the level of UNC8732 decreased below the detection limit as early as 4 hours post-treatment (Figure 2C). However, when UNC8732 was added to DMEM media without FBS, we did not observe the formation of the aldehyde metabolite, suggesting that amine oxidases in the FBS are responsible for this transformation (Figure 2D). When exposed to the same conditions, UNC8153 behaves similarly (Extended Data Figure 2A-B). Furthermore, the addition of aminoguanidine (AG), a pan-amine oxidase inhibitor, to UNC8732 in the presence of FBS-containing DMEM media effectively inhibits UNC8732 metabolism (Extended Data Figure 2C), supporting that amine oxidases in the FBS are responsible for this metabolic transformation. In the presence of FBS, the addition of AG inhibited NSD2 degradation by UNC8732 in a dose-dependent fashion (Figure 2E). Last, we treated U2OS cells with UNC8732 in the presence of 10% FBS or the absence of FBS. In the absence of FBS, no NSD2 degradation was observed (Figure 2F). These data strongly suggest that the metabolism of UNC8732 to the aldehyde is required for effective degradation.

We next sought to synthesize the aldehyde metabolites of UNC8153 and UNC8732; however, our initial efforts to access the aldehydes in high purity were unsuccessful. Instead, we synthesized and isolated the pure alpha-hydroxy sulfonates UNC9801 and UNC10088 (Figure 2G). These analogs, also known as aldehyde bisulfite adducts, are known hydrolytically labile prodrugs (or protecting groups) of aldehydes.^20^ While the amine-containing compounds UNC8153 and UNC8732 require enzymatic oxidation to yield the corresponding aldehydes, these alpha-hydroxy sulfonates convert via simple hydrolysis under aqueous conditions.

In order to confirm that the alpha-hydroxy sulfonates exhibit rapid transformation to the corresponding aldehydes, we first monitored this conversion under assay conditions (media +/- FBS). As anticipated, the aldehyde product is formed immediately upon exposure of UNC9801 to cell media, with the hydrolysis occurring so rapidly that the parent alpha-hydroxy sulfonate (UNC9801) is never detected (Extended Data Figure 2D). Next, we tested UNC9801 and UNC10088 in the ICW assay and demonstrated that they promote NSD2 degradation in a dose-dependent fashion (Figure 2H) (UNC9801 DC_50_ = 0.27 ± 0.15 µM and D_max_ = 87 ± 6%; UNC10088 DC_50_ = 0.06 ± 0.01 µM and D_max_ of 97 ± 3%). Interestingly, both UNC9801 and UNC10088 exhibit DC_50_ values that are comparable to their amine analogs, with UNC8732 and UNC10088 being more potent than UNC8153 and UNC9801 (Figure 2H). When evaluating NSD2 degradation in a time-dependent fashion, we found that UNC8153 and its matching bisulfite UNC9801 appear to have similar time-to-D_max_ at around 4-6 hours (Figure 2I). UNC8732 and UNC10088 are not only more potent, but they also degrade NSD2 more rapidly compared to UNC8153 and UNC9801, reaching D_max_ at around 2-4h (Figure 2I). After 0.5h, treatment with UNC10088 results in more significant NSD2 degradation as compared to UNC8732, potentially due to the more rapid conversion of UNC10088 to the active aldehyde. Last, we evaluated the ability of UNC10088 to degrade NSD2 in the absence of FBS and found that, unlike UNC8732, it does not depend on the presence of FBS to promote NSD2 degradation (Figure 2J). Importantly, we previously synthesized and tested the corresponding acid analog of UNC8153, which did not promote NSD2 degradation, ruling out the possibility that further oxidation of the aldehyde plays a role in the observed degradation^9^. Overall, these results confirm that our primary amine-containing NSD2 degraders are functioning as prodrugs and require metabolism to the corresponding aldehyde for activity.

### UNC8732 recruits the SCF^FBXO22^ ubiquitin ligase complex to promote NSD2 degradation

Next, we turned our attention towards identifying the putative E3 ubiquitin ligase that our compounds recruit to facilitate NSD2 degradation. We had previously shown that UNC8153 acts in a proteasome- and neddylation-dependent manner^9^ and confirmed that this is also the case for our second-generation degrader, UNC8732 (Extended Figure 3A-B). These data implicate one or more Cullin E3 ligases in our degradation mechanism. We carried out an unbiased, proteome-wide search for a candidate ubiquitin E3 ligase involved in degrading NSD2, using proximity-dependent biotin identification (BioID)^21^, a proximity biotinylation-based technique used to identify protein-protein interactions in living cells. Flp-In 293 cells expressing an NSD2-miniTurbo fusion protein (Figure 3A) were treated with either DMSO or UNC8732 in the presence of a proteasome inhibitor to prevent NSD2 degradation. A streptavidin pulldown of the resultant biotinylated proteins was analyzed by MS/MS (Figure 3B, Extended Data Figure 3C-E). Among the relatively few proteins significantly enriched upon UNC8732 treatment was FBXO22 (Figure 3B-C), an F-box containing substrate recognition subunit of the Neddylation-dependent SKP1-CUL1-F-box (SCF) E3 ligase complexes.^22,23^ SKP1 is an adaptor protein that bridges F-box proteins (such as FBXO22) to CUL1, and CUL1 recruits ubiquitin-charged E2 enzymes for ubiquitin transfer to substrates in a neddylation-dependent manner.^22,23^ Aside from FBXO22, there are also some increased interactions with 26S proteasome subunits, including PSMB1, PSMB5, and PSMA5 (Figure 3B).

**Figure 3:**
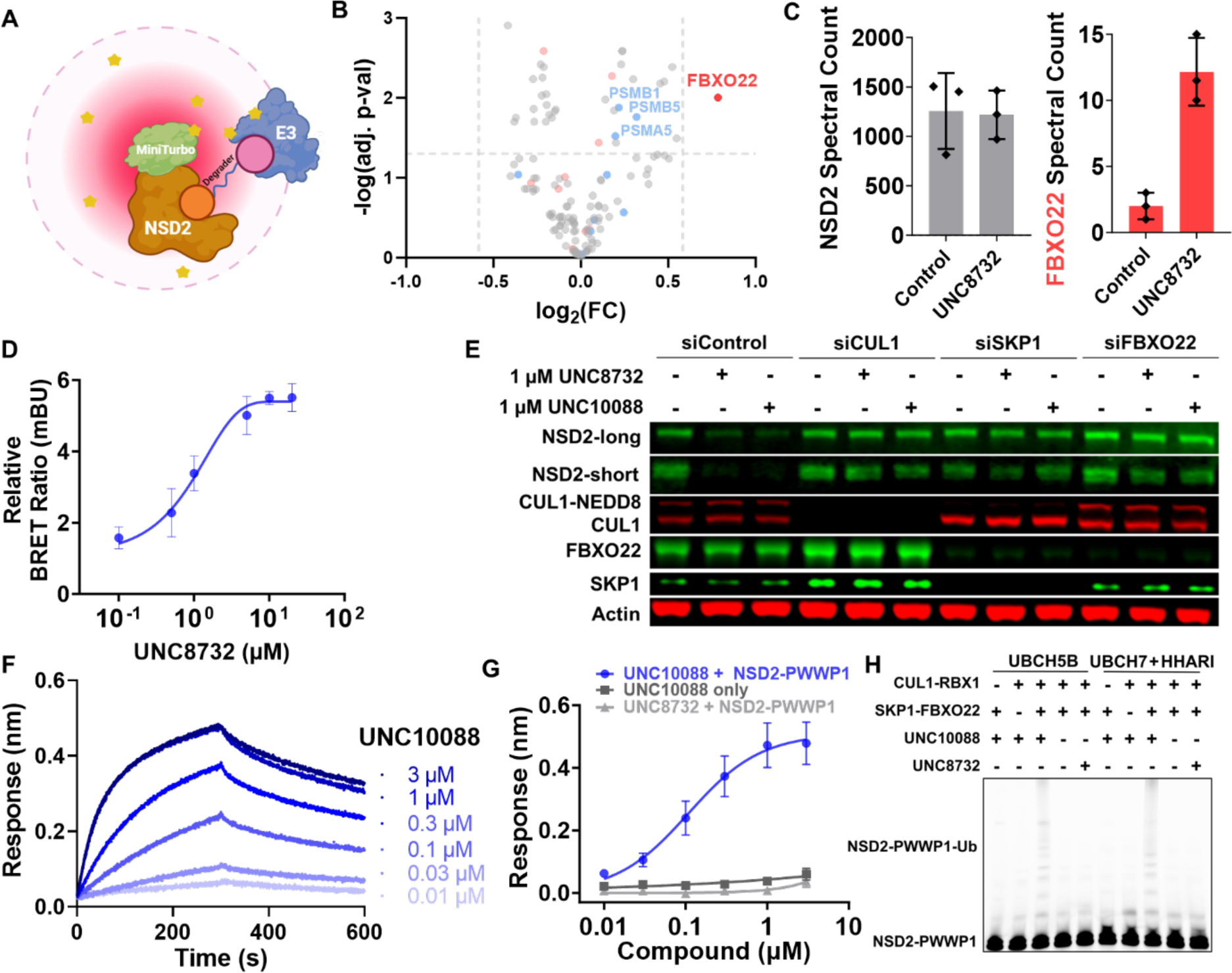
FBXO22 is responsible for compound-mediated degradation of NSD2. (A): Schematic showing a representation of the NSD2-miniTurbo fusion protein (brown and green), putative UNC8732 mediated interaction with an unknown E3 ubiquitin ligase (blue), and the approximate radius (10 nm) of biotinylation events^26^ generated by the NSD2-miniTurbo (pink). (B): Volcano plot of BioID results showing log_2_ fold change (FC) and -log_10_ adjusted p-values derived from normalized protein spectral counts. Proteins were previously filtered for SAINT BFDR ≤ 0.01 in either DMSO control or UNC8732 treated conditions. Red data points represent proteins in the GO:BP protein ubiquitination term (GO:0016567), and blue data points represent proteins included in the GO:CC proteasome core complex term (GO:0005839). Significance cut-off set at adj. p-value <0.05 and fold change >1.5. (C): Histogram showing the spectral counts of NSD2 and FBXO22. Data presented are the mean ± SD of 3 independent experiments. (D): A NanoBRET protein-protein interaction assay with NanoLuc-NSD2 and FBXO22-HT in U2OS cells reveals ternary complex formation. U2OS cells were treated with 0.05-40 µM of UNC8732 24h post-transfection for 3 hours. BRET ratios (mBU) are shown relative to DMSO treatment control (= 1.00). The data presented are the mean ± SD of 3 independent experiments. (E): Knockdown of SCF^FBXO22^ components by siRNA in U2OS cells. UNC8732 and UNC10088 degrader treatments were performed 4 days post-siRNA transfection for 4h. The immunoblot presented is representative of three independent experiments. (F): Representative BLI sensorgrams upon the addition of increasing concentrations of UNC10088 and a fixed concentration of NSD2-PWWP1 (2 μM). SKP1-FBXO22 was loaded on SA biosensors to an average response of 1 nm. Curves are shown as the average of three independent experiments. (G): Binding curve with the average BLI response calculated based on steady-state approximation. The data presented are the mean ± SEM of three independent experiments. (H): In vitro NSD2-PWWP1 ubiquitination with the indicated sets of substrate priming machinery, UBCH5B or UBCH7 with HHARI. Each reaction also contained CDC34B. The reaction mixture was analyzed by SDS-PAGE and fluorescence scanning. NSD2-PWWP1 was subject to a sortase reaction for fluorescent labeling. The gel presented is representative of three independent experiments.

To further investigate the putative UNC8732-mediated recruitment of FBXO22 to NSD2, we performed a proximity-based NanoBRET protein-protein interaction assay to assess the formation of an NSD2-degrader-FBXO22 ternary complex. We co-transfected U2OS cells with full-length NSD2 protein fused to the nanoluciferase protein (nanoLuc), and FBXO22 is fused to a HaloTag which covalently binds the HaloLigand (Promega 618 ligand)^24,25^. Bioluminescence resonance energy transfer (BRET) was observed between the two fusion proteins in a dose-dependent manner in the presence of increasing concentrations of UNC8732, indicating that these two proteins are brought into proximity upon degrader treatment in cells (Figure 3D).

Finally, to confirm the dependency of NSD2 degradation on the recruitment of FBXO22, an siRNA knockdown study was performed. The knockdown of any of the core components of the SCF^FBXO22^ complex, including CUL1, SKP1, or FBXO22, was able to rescue NSD2 levels and prevent UNC8732 and UNC10088 mediated NSD2 degradation (Figure 3E). These data confirm that FBXO22 and the SCF complex are required for NSD2 degradation mediated by UNC8732 and UNC10088. Overall, our data strongly support SCF^FBXO22^ as the UNC8732-recruited E3 ubiquitin ligase complex.^9^

### UNC10088 promotes ternary complex formation and NSD2 polyubiquitination *in vitro*

To further characterize the mechanism of action of our degraders, we performed several in vitro assays using UNC10088, which rapidly hydrolyzes to the corresponding aldehyde in buffer, and a recombinant SKP1-FBXO22 fusion protein with a C-terminal biotinylated Avi-Tag (Extended Data Figure 3F-G). A bio-layer interferometry (BLI) assay was utilized in which SKP1-FBXO22 was loaded onto streptavidin biosensors and evaluated for interaction with a fixed concentration of recombinant NSD2-PWWP1^14^ in the presence or absence of UNC10088. In the absence of UNC10088, no detectable binding signal was observed; however, a strong dose-dependent increase in NSD2-PWWP1 binding was observed with increasing concentrations of UNC10088 (Figure 3F-G), demonstrating the formation of a ternary complex. As expected, no ternary complex formation was observed upon treatment with UNC8732 (Figure 3G), which is unable to convert to its corresponding aldehyde in a buffer lacking an amine oxidase. This further supports the aldehyde species as the critical mediator of NSD2 degradation. Moreover, size exclusion chromatography demonstrated co-elution of SKP1-FBXO22 with NSD2-PWWP1 in the presence of UNC10088, providing further support for ternary complex formation in vitro (Extended Data Figure 3H). We used a stable SKP1/FBXO22 complex co-expressed from Sf9 insect cells and performed a similar experiment with consistent results (Extended Data Figure 3I).

Next, we aimed to demonstrate FBXO22-mediated NSD2 polyubiquitination in vitro. Specifically, we incubated recombinant CUL1, SKP1-FBXO22 fusion protein, NSD2-PWWP1, and the E2 ubiquitin-conjugating enzymes UBCH5B or UBCH7/HHARI with UNC8732 or UNC10088 and evaluated polyubiquitination of NSD2-PWWP1 by SDS-PAGE. In this experiment, polyubiquitination of NSD2-PWWP1 was observed only in the presence of all required components of the E3 ubiquitin ligase complex, including FBXO22 and UNC10088 (Figure 3H). In contrast, UNC8732 was not able to facilitate polyubiquitination of NSD2-PWWP1 in the *in vitro* buffer setting. These results clearly demonstrate that the FBXO22-NSD2 ternary complex mediated by hydrolyzed UNC10088 results in the polyubiquitination of NSD2 and helps to rule out other possible degradation mechanisms.

### Aldehyde degraders are dependent on cysteine 326 in the FIST_C domain of FBXO22

To further understand the mechanism of action of our NSD2 degraders, we investigated the specific nature of their interaction with FBXO22. First, we were interested in evaluating if our compounds would bind directly to FBXO22, independent of NSD2-PWWP1. We synthesized a truncated analog of UNC8732 containing the putative E3 recruiting end of the molecule (UNC9630), as well as an inactive methylated amine analog (UNC9631, Figure 4A). We previously showed that methylation of the terminal amine of UNC8153 results in loss of NSD2 degradation, presumably due to an inability of the methyl amine to be oxidized to an aldehyde.^9^ Both truncated compounds were tested for their ability to compete with UNC8732-mediated NSD2 degradation in the ICW assay (Figure 4A). While the addition of UNC9630 effectively competes with UNC8732 for degradation, with NSD2 levels increasing with increasing concentrations of UNC9630, UNC9631 has no effect even at the highest concentrations. This reveals that the aldehyde metabolite of UNC9630 is sufficient to engage FBXO22. To further evaluate FBXO22 binding, we performed differential scanning fluorimetry (DSF) experiments and observed a dose-dependent thermal stabilization of the co-expressed SKP1/FBXO22 complex (as used in Extended Data Figure 3I) with increasing concentrations of UNC10088 (Figure 4B-C). Consistent with the BLI results, the primary amine-containing UNC8732 did not stabilize SKP1/FBXO22 in the same way.

**Figure 4:**
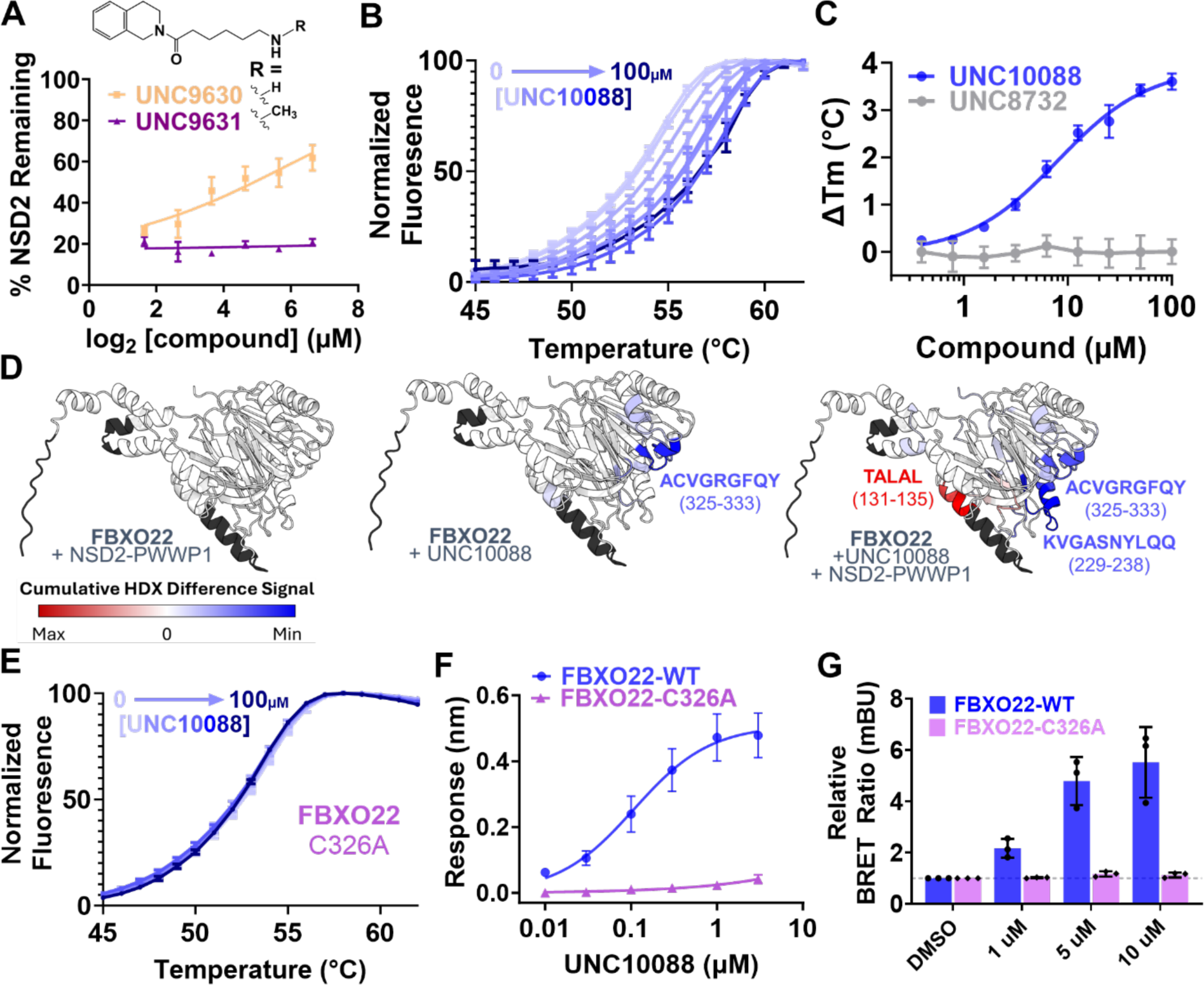
Aldehyde degraders bind FBXO22 in a cysteine 326-dependent manner. (A): In-cell western quantification of NSD2 levels in U2OS cells co-treated with 1 µM of UNC8732 and 6.25-100 µM of UNC9360 (E3-recruiting handle of UNC8732) or UNC9631 (monomethylated E3-recruiting handle of UNC8732) for 6h. Data represents the average ± SEM of 4 technical replicates from 2 independent experiments. (B): Normalized fluorescence data from DSF upon incubation of co-expressed SKP1/FBXO22 with 0-100 µM UNC10088. Data represents the average ± SD of 3 independent experiments. (C): Change in melting temperature (Tm) as measured by DSF of co-expressed SKP1/FBXO22 treated with 0-100 µM UNC8732 and UNC10088. Data represents the average ±SD of 3 independent experiments. (D): AlphaFold predicted structure of FBXO22 highlighted to indicate the cumulative HDX difference signal in an HDX-MS experiment. Blue = reduced uptake, red = increased uptake, black = no FBXO22 sequence coverage. For clarity, SKP1 is not shown in the AlphaFold structure, and no changes in deuterium uptake were observed for SKP1 (Extended Data Figure 4B). (E): Normalized fluorescence data from DSF upon incubation of co-expressed SKP1/FBXO22 with 0.1-100 µM UNC10088. Error bars represent the SD of 4 technical replicates from one independent experiment. (F): Binding curve of the average BLI response for both SKP1-FBXO22 WT and C326A calculated based on steady-state approximation. Data represents the average ± SEM of three independent experiments. (G): NanoBRET assay results with NanoLuc-NSD2 and FBXO22-HT wildtype or C326A mutant performed in U2OS cells. U2OS cells were treated with the indicated concentration of UNC8732 24h post-transfection for 3 hours. BRET ratios (mBU) are shown relative to DMSO treatment control (= 1.00). Data represents the average ± SD of 3 independent experiments.

Given the small size and simplicity of the alkyl aldehyde, we were intrigued as to how this group may be binding and recruiting FBXO22 on a molecular level. As aldehydes are mildly electrophilic species and are capable of covalent reaction with nucleophilic amino acid side chains such as Lys or Cys in an irreversible or reversible fashion, respectively, we hypothesized that the aldehyde species might be reacting covalently with a nucleophilic residue on FBXO22. To investigate where on FBXO22 our compounds may be binding, we used Hydrogen Deuterium Exchange-Mass Spectrometry (HDX-MS). In this experiment, we used the stable SKP1/FBXO22 complex (as used in Extended Data Figure 3I) and evaluated three different conditions: SKP1/FBXO22 + NSD2-PWWP1, SKP1/FBXO22 + UNC10088, and SKP1/FBXO22 + UNC10088 + NSD2-PWWP1. In each case, the relative deuterium uptake (defined as the number of deuterium exchanged divided by the number of exchangeable backbone amides of a given peptide) was compared to that of the SKP1/FBXO22 apo protein complex, represented by a differential HDX-MS signal (ΔHDX-MS, %). The expectation is that ΔHDX-MS would yield non-zero values in regions of ligand binding, protein-protein interactions, or as a result of allosteric changes.^27^ Stabilization of the protein backbone, causing a persistence of hydrogen bonding and reduced deuterium uptake, is attributed to peptides at binding or protein-protein interfaces and results in a negative ΔHDX-MS signal.^28^ To be considered statistically significant, the cumulative (sum of all timepoints tested) ΔHDX-MS signal must have exceeded the cumulative error by > 1%. Unsurprisingly, there were no significant ΔHDX-MS signals upon incubation of SKP1/FBXO22 with NSD2-PWWP1 in the absence of UNC10088 (Figure 4D left, Extended Data Figure 4A), in agreement with our BLI results (Figure 3G-H). In contrast, the addition of UNC10088 to SKP1/FBXO22 caused a significant reduction in deuterium uptake in a peptide corresponding to FBXO22 residues 325-333 (Figure 4D, middle). These residues are in the FIST_C domain of FBXO22, a domain unique to FBXO22 and the only domain of this kind in the human proteome.^29^ Within this region of reduced deuterium uptake, cystine 326 provoked our interest due to its potential nucleophilicity. HDX measurements for the 3-component system, including SKP1/FBXO22, UNC10088, and NSD2-PWWP1, also showed a reduction in deuterium uptake in FBXO22 residues 325-333, with an additional region protected from deuterium uptake at residues 229-238. When mapped onto the AlphaFold predicted structure of FBXO22, residues 229-238 are on the same side of the FIST_C domain as residues 325-333 (Figure 4D right), suggesting that the newly protected residues may engage NSD2-PWWP1 when the compound induces a ternary complex. Interestingly, an increase in deuterium uptake was observed for residues 131-135, which map to the opposite side of the putative FBXO22-degrader-NSD2 binding interface, suggesting a potential allosteric change in protein structure when the ternary complex is formed.

To better understand the role of C326 in the recruitment of FBXO22 by our degraders, we generated a C326 to alanine mutation and found that it abolished the interaction with UNC10088. Specifically, UNC10088 did not influence the melting temperature of the C326A mutant SKP1/FBXO22 complex as determined by DSF (Figure 4E). Importantly, the C326A mutation did not significantly destabilize the melting temperature of the recombinant domain, suggesting that the structural integrity of the mutant protein was preserved. Similarly, the introduction of the C326A mutation in the recombinant SKP1-FBXO22 fusion protein prevented ternary complex formation with NSD2-PWWP1 and UNC10088 *in vitro* as determined by BLI (Figure 4F; Extended Data Figure 4C) and *in vitro* polyubiquitination of NSD2 could not be achieved in the presence of C326A-mutant SKP1-FBXO22 (Extended Data Figure 4D). Finally, the C326A mutation in FBXO22 abolished the ability of UNC8732 to induce ternary complex formation in the cellular NanoBRET assay (Figure 4G) with no difference in the expression level of WT compared to the mutant FBXO22-HT plasmid observed (Extended Data Figure 4E). Together, this data strongly suggests that the aldehyde derivative of UNC8732 and UNC10088 binds and recruits FBXO22 via interactions with C326.

### FBXO22 binding with UNC10088-aldehyde is likely a reversible covalent interaction

Given the reactive nature of cystine residues, we hypothesized that the aldehyde derivative of UNC10088 is likely forming a reversible covalent bond with C326 via the formation of a hemithioacetal (Extended Data Figure 4H). To test the reversible nature of this interaction, we performed a washout experiment and monitored the thermal stability of SKP1/FBXO22 by DSF (Extended Data Figure 4F). Specifically, we incubated SKP1/FBXO22 and UNC10088 for 20 minutes, followed by three successive dilutions and filtration/reconcentration through a 50 kDa molecular weight cutoff membrane to washout any available compound. The final ‘washed’ sample was evaluated by DSF relative to SKP1/FBXO22 apo and UNC10088-bound proteins. As expected, an increase in thermal stabilization was observed for SKP1/FBXO22 in the presence of UNC10088 (Extended Data Figure 4G). However, upon dilution and reconcentration, the melting temperature reverted to that of the apo state (Extended Data Figure 4G). This reduced stability of the ‘washed out’ sample supports a covalent *reversible* interaction between the aldehyde adduct of UNC10088 and C326 of FBXO22 (Extended Data Figure 4H). Moreover, the reversible nature is further supported by the steady MS signal intensity (ion abundance) for FBXO22 peptide 325-333 in the presence and absence of UNC10088.

### Alkyl amine containing XIAP degrader recruits the SCF^FBXO22^ complex for XIAP degradation

We hypothesized that the primary amine FBXO22 warhead may hold potential for use more broadly to hijack FBXO22 for TPD. To this end, we were intrigued by the recently reported degrader of XIAP containing a primary alkyl amine as its functional degradation-inducing moiety (Figure 5A)^13^ and we sought to further investigate the mechanism of action of the XIAP degrader (Compound 10) to understand whether its degradation mechanism was related to that of our NSD2 degraders.

**Figure 5:**
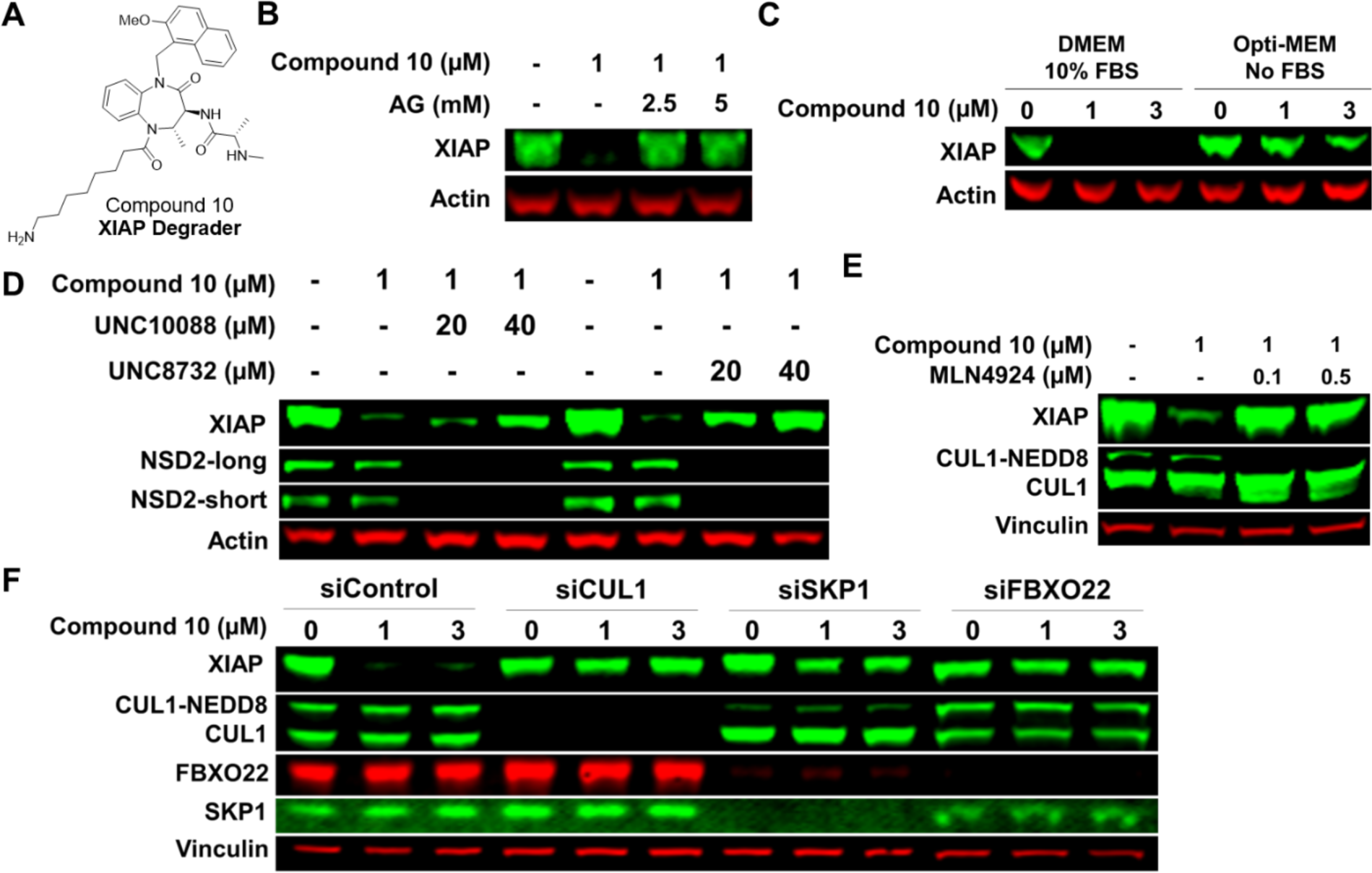
A primary amine-containing degrader recruits the SCF^FBXO22^ complex for degradation of XIAP. (A): Chemical structure of primary amine-containing XIAP degrader, Compound 10, reported by den Besten et al.^13^ (B): MCF7 cells were treated with Compound 10 and varying concentrations of aminoguanidine (AG) for 24 hours, followed by immunoblotting analysis for XIAP levels. Actin is a loading control. (C): MCF7 cells were cultured in DMEM + 10% FBS or Opti-MEM with no FBS and subsequently treated with the indicated concentrations of Compound 10 for 24h, followed by immunoblotting analysis for XIAP levels. Actin is a loading control. (D): MCF7 cells were treated with Compound 10 and varying concentrations of UNC10088 and UNC8732 for 24 hours, followed by immunoblotting for XIAP and NSD2. Actin is a loading control. (E): MCF7 cells were treated with Compound 10 and varying concentrations of MLN4924 (a selective neddylation inhibitor ^30^) for 24 hours, followed by immunoblotting for XIAP levels. CUL1 was blotted as a representative Cullin. Vinculin is a loading control. (F): siRNA knockdown of core SCF^FBXO22^ components in MCF7 cells was performed. Compound 10 treatments were performed 72h post-siRNA transfection for 24h. Vinculin is a loading control.

First, we investigated whether XIAP degradation is dependent on oxidation of the primary amine of Compound 10. We found that the addition of the amine oxidase inhibitor aminoguanidine (AG) was similarly able to inhibit the degradation of XIAP mediated by Compound 10 (Figure 5B). Additionally, removal of FBS from the culture medium prevented Compound 10-induced XIAP degradation (Figure 5C), similar to the FBS dependence observed with our NSD2 degrader. We next performed a competition experiment, co-treating cells with the XIAP degrader and our NSD2 degraders, UNC8732 and UNC10088. The presence of excess UNC10088 or UNC8732 significantly reduced the degradation efficiency of the XIAP degrader, suggesting competition between the degraders for recruitment of the same E3 ligase. Further, we demonstrated that XIAP degradation is also neddylation dependent (Figure 5E) in addition to being proteasome dependent^9^, suggesting that the XIAP degrader recruits a Cullin complex to mediate degradation. Finally, to test the possible role of the SCF^FBXO22^ complex in Compound 10 mediated XIAP degradation, we evaluated XIAP degradation upon knockdown of CUL1, SKP1, and FBXO22. The knockdown of each complex member resulted in the inhibition of XIAP degradation by Compound 10 (Figure 5F). These findings strongly suggest that the previously reported Compound 10 is also able to recruit FBXO22 to facilitate XIAP degradation and provides evidence that the FBXO22 recruiting mechanism by a primary amine prodrug may have applicability beyond NSD2 targeted degradation.

## Discussion

In this work, we report the discovery of highly potent degraders of NSD2, which importantly revealed a novel approach to achieve targeted protein degradation through the recruitment of FBXO22. We developed two types of prodrug degraders, both yielding the same active aldehyde species, which can promote NSD2 degradation. The primary amine-containing compound, UNC8732, is oxidized through the activity of an amine oxidase(s) in FBS and is well-suited for cellular studies. Indeed, the use of UNC8732 to probe the consequence of NSD2 degradation in gain-of-function NSD2 mutant ALL cell lines revealed an exciting and potentially therapeutically relevant phenotype. In addition to the induction of apoptosis and reduction of cell viability, UNC8732 was able to overcome the resistance of the E1099K NSD2 ALL cells to glucocorticoid therapy, an important clinical problem in relapsed ALL.^16^ Given these results, UNC8732 can serve as a valuable chemical probe to study both fundamental aspects of NSD2’s function in nuclear signaling and epigenetics, as well as the therapeutic potential of targeting NSD2 for degradation. UNC8884 is a structurally similar compound that does not bind to NSD2 due to lack of a key cyclopropyl group^14^ and, therefore, does not degrade NSD2, providing an excellent inactive control for use alongside UNC8732. The corresponding alpha-hydroxy sulfonate of UNC8732, UNC10088, functions as a prodrug through a simple hydrolysis reaction to form an aldehyde and is well suited for *in vitro* biochemical or biophysical assays in buffers lacking oxidase activity.

UNC8732 recruits FBXO22, a subunit of a SKP1, CUL1, F-box complex (SCF^FBXO22^) to promote NSD2 degradation. BLI, SEC, HDX-MS, and NanoBRET provide strong evidence of ternary complex formation between the NSD2 PWWP1 domain and FBXO22 upon the addition of UNC8732 and/or UNC10088. We show that the aldehyde moiety of our degraders likely interacts via a reversible covalent interaction with Cys326 of the FBXO22 FIST_C domain, as mutation of Cys326 abolishes ternary complex formation. The FIST_C domain is a substrate-targeting domain of FBXO22, while the F-box region interacts with SKP1, suggesting that degrader-mediated ternary complex formation via the FIST_C domain could catalyze polyubiquitylation. Indeed, UNC10088 promotes ubiquitylation of NSD2-PWWP1 in a Cys326-dependent manner.

We have further shown that a previously reported primary amine-containing degrader of XIAP (Compound 10)^13^ is also dependent on the SCF^FBXO22^ E3 ligase complex for its degradation activity. Similar to UNC8732, the activity of Compound 10 is blocked in the presence of an amine oxidase inhibitor, suggesting that it may also function through a similar aldehyde-mediated interaction with FBXO22. Compound 10 was proposed to act through a mechanism in which XIAP – itself an E3 ligase – ubiquitylates compound 10 at the latter’s primary amine group, leading to polyubiquitylation of the compound, which, when bound to XIAP could promote proteasomal degradation. Our data suggests that a more classical TPD mechanism involving the recruitment of SCF^FBXO22^ to XIAP by Compound 10 is also possible. These data suggest that FBXO22-mediated degradation is not necessarily specific to NSD2 and supports the potential use of simple primary amine warheads to recruit FBXO22 for TPD in the future.

FBXO22 has been reported as an E3 ligase responsible for the degradation of several proteins, such as the proapoptotic transcription factor BACH1^33^, the cell cycle inhibitor p57^Kip2^,^34^ and the nuclear fraction of tumor suppressor PTEN^35^, among others.^31^ Thus, FBXO22 is indeed capable of degrading nuclear proteins such as NSD2. While FBXO22 is expressed in most cell types ^31^ and may, therefore, be an attractive E3 ligase for broad use in TPD, it has also been shown to have elevated expression in various types of human tumor tissues compared to that in normal tissues, further supporting the use of this TPD strategy for applications in oncology.^31,32^ Overall, we believe there is much potential for this novel FBXO22-dependent TPD strategy to broaden the scope of available TPD reagents and therapeutics and for UNC8732 as a tool to explore NSD2 disease phenotypes.

## Acknowledgments

The authors thank the members of the James laboratory and Stephen Frye for helpful discussions and input throughout the project. The authors thank Peter H. Buttery and Juanita L. R. Sanchez for the review of experimental data. This work is supported by grants from the Canadian Institutes of Health Research (FDN154328, <u>OGB</u>190363) and the Princess Margaret Cancer Foundation to C.H.A., from the National Institutes of Health (NCI) (R01CA242305) to L.I.J., and a Leukemia and Lymphoma Society Specialized Center for Research and Florida Department of Health Grant 22L03 to J.D.L. D.Y.N. is supported by a Canadian Graduate Scholarship – Doctoral Research Award from the Canadian Institutes of Health Research (494204) and a Doctoral Training Scholarship from Fonds de recherche du Québec - Santé (320128). J.R.T. is supported by the UNC Lineberger Comprehensive Cancer Center Cancer Epigenetics Training Program (5T32CA217824-05). D.B-L. is supported by CRS grant 25418. D.W. is supported by the NSERC Discovery (RGPIN-480432) and NSERC Collaborative Research and Development (CRDPJ-504037) grants. Portions of this work have been supported by certain funds managed by Deerfield Management Company, L.P. Deerfield Management Company is a healthcare-focused investment management firm. The Structural Genomics Consortium is a registered charity (No. 1097737) that receives funds from Bayer AG, Boehringer Ingelheim, BristolMyersSquibb, Genentech, Genome Canada through Ontario Genomics Institute [OGI-196], EU/EFPIA/OICR/McGill/KTH/Diamond Innovative Medicines Initiative 2 Joint Undertaking [EUbOPENGrant875510], Janssen, Merck KGaA (aka EMD in Canada and US), Pfizer and Takeda. This material is based in part upon work supported by the National Science Foundation under Grant No. CHE-1726291. The authors thank the University of North Carolina’s Department of Chemistry Mass Spectrometry Core Laboratory for their assistance with mass spectrometry analysis. The authors thank Syngene International Ltd. for support of portions of this work.

## Methods

### In-Cell Western Assay (ICW)

U2OS cells were seeded to achieve about 80% confluency in Nunc™ MicroWell™ 96-Well Flat Clear Bottom Black Microplate (ThermoFisher 165305) for 96-well format or Corning® 384-well Flat Clear Bottom Black plates (Corning 3764) for 384-well format. Compounds were diluted in DMEM medium and added to the cells to achieve the prescribed final concentrations.

At the treatment endpoint, the medium was removed completely from all wells, and each well was rinsed very gently with PBS once. The cells were fixed in each well with 2% formaldehyde (37.5% stock from Sigma-Aldrich diluted in 1x PBS; 15 μL for 384-well or 50 μL for 96-well) for 10 minutes at room temperature. The wells were washed 3x with PBS and were then permeabilized with 0.25% Triton X-100 (diluted in 1x PBS; 15 μL for 384-well or 50 μL for 96-well) for 15 min at room temperature. It was subsequently washed 3x with PBS and then blocked by incubating all wells with 5% Bovine serum albumin (BSA) (diluted in 1x PBS-T; 15 μL for 384-well or 50 μL for 96-well) for 1 hour at room temperature or at 4 ᵒC overnight, followed by incubation with NSD2 primary antibody (Abcam ab75359; RRID:AB_1310816) diluted at 1:1000 in 5% BSA (at 15 μL for 384-well or 50 μL for 96-well) for 1 hour at room temperature with agitation. At least two wells were intentionally not incubated with NSD2 primary antibody (for background subtraction). Following NSD2 antibody incubation, the cells were washed 3x by PBS, then stained by LI-COR secondary antibody IRDye® 800CW Donkey anti-Mouse (LI-COR 926-32212; RRID:AB_621847) diluted 1:1000 and DRAQ5 (Cell Signaling #4084) diluted 1:2000 in 5% BSA for 1 hour at room temperature with agitation. The cells were washed 3x by PBS and imaged on a LI-COR ODYSSEY CLx near-IR scanner, and intensities of the 800CW and 700CW channels were quantified.

For data analysis, the 800CW intensity (NSD2) was normalized to 700CW intensity (DRAQ5), and the no NSD2 antibody control (background) normalized intensity was subtracted from the normalized intensity of each well. The background-subtracted normalized intensity was expressed as a percentage of the average background-subtracted normalized intensity of DMSO control treated wells, resulting in the numeric values for % NSD2 Remaining for each well. For each set of dose-response, the % NSD2 Remaining was plotted against the log of concentration, and the curves were fitted by [inhibitor] vs. response non-linear fit model in GraphPad Prism 10.1 with the Top constrained = 100 and the Bottom constrained = lowest % NSD2 Remaining for the dilution series. The DC_50_ was extracted from the calculated ‘IC50’ value of the curve-fitting, and the D_max_ was extracted from 100% – (lowest % NSD2 Remaining) for the dilution series.

### Surface Plasmon Resonance (SPR)

We conducted SPR experiments using a Biacore^TM^ 8K instrument at a temperature of 20°C, following previously established procedures^9,14^ with some adjustments. To begin, we immobilized the biotinylated NSD2-PWWP1 domain onto a CM5 chip (Cytiva) coupled with neutravidin, yielding approximately 1,500 response units (RU). The immobilization was carried out in HBS-EP+ buffer, which consist of 10 mM HEPES at pH 7.4, 150 mM NaCl, 3 mM EDTA, and 0.05% Tween 20. Compounds were tested at the highest concentration of 2 μM and diluted to a 1:3 ratio to generate five different concentration points. The experiments were conducted in the same buffer, supplemented with 0.5% DMSO in a single cycle kinetics with a 60s association time and a subsequent dissociation time of 120 s, all at a 40 µl/min flow rate. Data analysis was performed using the Biacore^TM^ Insight Evaluation software with blank and reference subtraction.

### NSD2 Degrader and Dexamethasone Treatment

Isogenic acute lymphoblastic leukemia (ALL) cell line RCH-ACV harboring the NSD2 p.E1099K mutation or edited to express wild type (WT) NSD2^19^ were cultured in RPMI-1640 medium with 10% FBS and treated with varying concentrations of NSD2 degrader (UNC8732) and negative control (UNC8884) for up to 21 days. The cells were then divided into two groups at each time point. One group was continuously treated with NSD2 degrader, and the other was treated with NSD2 degrader and dexamethasone (Sigma-Aldrich, #D9184) (1 µM) for 72 hours. The cells were harvested for cell viability assay, apoptosis, and immunoblotting.

### Cell Viability Assay

The cell viability of isogenic RCH-ACV cell lines treated with NSD2 degrader and dexamethasone was determined by the CellTiter-Glo® Luminescent Cell Viability Assay (Promega, #G7572). The opaque-walled 96-well plates with cells were equilibrated at room temperature before detection. Equal volumes of CellTiter-Glo Reagent to the volume of cell culture medium were added into each well and mixed for 2 minutes on a shaker to induce cell lysis. The plates were incubated at room temperature for 10 minutes to stabilize the luminescent signal before measurement of cellular viability using CLARIOstar (BMG LABTECH).

### Detection of apoptosis using Annexin V/PI staining

Isogenic RCH-ACV cell lines treated with NSD2 degrader and dexamethasone were harvested for apoptosis detection using Annexin V FITC Apoptosis Detection Kit (eBioscience, #BMS500FI-300). The cells were washed with PBS, resuspended in 1X Binding Buffer and incubated with 5 µL of FITC Annexin V for 10 min and 10 µL of Propidium Iodide (PI). Stained cells were required for apoptosis analysis by flow cytometry BD LSRFortessa™ (RRID:SCR_019600) within one hour. The data was analyzed by FlowJo_v10.7.1.

### RCH-ACV Cell Lysate Immunoblotting

After treatment of NSD2 degrader or dexamethasone, isogenic RCH-ACV cell lines were harvested for protein extraction using Minute^TM^ Total Protein Extraction Kit (Invent Biotechnologies, #SD-001/SN-002). Proteins were separated using SDS-PAGE and transferred to nitrocellulose membranes using iBlot 2 (Invitrogen). After blocking for one hour, the membrane was incubated with primary antibodies including anti-NSD2 (Abcam, RRID:AB_1310816), anti-HDAC2 (Millipore, RRID:AB_11212215), anti-H3K36me2 (Abcam, RRID:AB_1280939), anti-H3K27me3 (Cell Signaling, RRID:AB_2616029), anti-H3 (Cell Signaling, RRID:AB_2756816) at 4°C overnight. The membranes were then incubated with second antibodies including Alexa Fluor™ Plus 680 Goat anti-Rabbit IgG (H+L) (Invitrogen, RRID:AB_2633283) or Alexa Fluor™ Plus 800 Goat anti-Mouse IgG (H+L) (Invitrogen, RRID:AB_2633279) for one hour. After washing three times with PBST, the membranes were scanned using an Li-Cor Odyssey CLx Imaging System (RRID:SCR_014579).

### U2OS and MCF7 Cell Culture

U2OS and MCF7 Cell lines were cultured in DMEM supplemented with 10% fetal bovine serum (FBS) (Wisent) and 100 U/ml penicillin and 100 μg/ml streptomycin (Wisent) at 5% CO_2_ at 37 °C in accordance with standard mammalian tissue culture protocols and periodically tested for mycoplasma contamination via the MycoAlert Mycoplasma Detection kit (Lonza).

### Cell Treatment in Opti-MEM

500 µL of 2 × 10^5^ cells/mL were seeded into 24-well plates in DMEM medium with 10% FBS overnight for cell attachment. Upon cell attachment, the DMEM medium was completely removed, and the wells were washed gently with 1 mL PBS twice to get rid of any remaining residue of DMEM before adding 500 µL of Opti-MEM (Gibco 31985070) with no FBS. The compounds were subsequently added to cells cultured in Opti-MEM and allowed to incubate for the prescribed treatment time.

### MLN4924, MG-132, Aminoguanidine (AG) Treatments

500 µL of 2 × 10^5^ cells/mL were seeded into 24-well plates in DMEM medium with 10% FBS overnight for cell attachment. The cells were treated with either MLN4924 (Cayman Chemical 15217, dissolved to 20 mM stock in DMSO), MG-132 (Abcam ab141003, dissolved to 20 mM stock in DMSO), or AG (Sigma-Aldrich 396494, dissolved to 500 mM stock in DMEM, made freshly for each experiment) to their prescribed concentrations. The cells were subsequently immediately treated with the degrader compound at the prescribed concentrations and incubated at cell culture conditions for the prescribed treatment time.

### U2OS and MCF-7 Cell Lysate Immunoblotting

Cell pellets were lysed directly in the culture plate or in 1 mL tubes for 3 minutes at room temperature with lysis buffer containing 20 mM Tris-HCl pH 8, 150 mM NaCl, 10 mM MgCl2, 1 mM EDTA, 0.5% Triton X-100, Protease Inhibitor (Roche 11873580001), and benzonase (Millipore Sigma E1014). Subsequently, SDS was added to achieve a final concentration of 1%. Next, the mixtures were spun on a tabletop centrifuge at maximum RPM for 10 minutes, and the supernatants (cell lysates) were transferred to a new tube. The concentration of protein in each lysate was measured by Thermo Scientific Pierce™ BCA Protein Assay Kit (Cat. # 23225) per supplier protocol. 10-30 mg of protein per loading was prepared in a sample loading buffer. Samples were loaded into Invitrogen NuPAGE Bis-Tris Gels and ran in 1X NuPAGE MOPS buffer (Invitrogen NP0001) for 90 minutes at 120 V. The gel was subsequently transferred by Bio-Rad Mini Trans-Blot® Cell at 70V/500mA for 1.5 h, per manufacturer protocol, in Tris-Glycine transfer buffer (3g/L Tris, 14.4g/L glycine and 10% methanol), on ice. Membranes were blocked in 5% BSA in PBS-T for 1 hour at room temperature, then blotted overnight by primary antibodies: NSD2 (1:1000; Abcam ab75359; RRID:AB_1310816), SKP1 (1:1000 dilution; Proteintech 10990-2-AP; RRID:AB_2187492), CUL1 (1:1000 dilution; Proteintech 12895-1-AP; RRID:AB_2086291), FBXO22 (1:1000 dilution; Proteintech 13606-1-AP; RRID:AB_2104403), XIAP (1:1000 dilution; Proteintech 66800-1-Ig; RRID:AB_2882143), Actin (1:5000 dilution; ab179467; RRID:AB_2737344) or Vinculin (1:2000; Santa Cruz sc-73614; RRID:AB_1131294) (1:2000; Cell Signaling 13901S; RRID:AB_2728768), diluted in 5% BSA. Upon primary antibody blotting, the membrane was washed for 5-min by PBS-T three times and subsequently blotted by LI-COR secondary antibodies: IRDye® 800CW Goat anti-Rabbit (LI-COR 926-32211; RRID:AB_621843), 680RD Donkey anti-Rabbit (926-68073; RRID:AB_10954442), 800CW Donkey anti-Mouse (926-32212; RRID:AB_621847), or 680RD Goat anti-Mouse (926-68070; RRID:AB_10956588) diluted 1:5000 in 5% BSA, for 1-hour at room temperature and washed for 5-min by PBS-T three times. The membrane was imaged on a LI-COR Odyssey CLx scanner.

### Metabolite identification for UNC8153 performed in U2OS cells in DMEM (with 10% FBS) incubations at 10 µM

A working stock solution of UNC8153 was prepared by adding 2 µL of 10 mM stock in DMSO to 10% (V/V) FBS (Gibelco)/DMEM (Gibelco; 998 µL). The working solution of UNC8153 (100 µL, 20 µM) was added to U2OS cell suspension (100 µL) in a 96-well plate. This cell suspension was prepared by seeding U2OS cells (ATCC; density 8000 cells/well) and incubating under 5% CO_2_ for 24 h at 37 °C. The samples (each containing 200 µL) were mixed with 600 µL of acetonitrile spiked with the internal standard tolbutamide (500 ng/mL) at each time point; then, each sample was vortexed and centrifuged at 4000 rpm for 15 minutes. After centrifugation, each supernatant was analyzed by LC-HRMS/MS (AB Sciex Triple TOF 5600+ mass spectrometer; Agilent 1290 Infinity II HPLC, GL Sciences, Inertsil C18-ODS, 3.0 × 150 mm, 5 µm column, solvent gradient 0.1% Formic acid in 5 mM ammonium formate/acetonitrile; 400 µL/min).

### Stability of UNC8732 in Culture Medium

The 2 µM working solution of the UNC8732 stock was added to DMEM or DMEM + 10% FBS (V/V) or DMEM + 10% FBS (V/V) + 10mM aminoguanidine such that the final concentration of the UNC8732 was 1 µM. The mixture was incubated at 37 °C. At specific time points, aliquots (50 µL) of the mixture were treated with acetonitrile (400 µL) containing 500 ng/mL tolbutamide as internal standard, then vortexed for 5 min. The suspension was centrifuged at 12000 rpm at 4 °C for 10 min, and the supernatant was analyzed by LC-MS/MS (AB Sciex 4500 QQQ MS system connected with Shimadzu Nexera X2 LC system). All samples were qualitatively monitored for UNC8732 and the aldehyde metabolite. Data are reported in area ratio (ratio of UNC8732 and aldehyde metabolite to Internal standard MS area response) at each time point for all samples.

### Stability of UNC9801 in Culture Medium

UNC9801 was diluted to 100 µM final concentration from 10 mM DMSO stock in DMEM and DMEM/10% FBS (v/v) by diluting 10 uL of the 10 mM DMSO stock. The mixture was incubated at 37 °C. After 0, 1, 4, 6, 12, and 24 h, aliquots (50 µL) of the mixture were treated with acetonitrile (400 µL) containing 500 ng/mL tolbutamide as internal standard, then vortexed for 5 min. The suspension was centrifuged at 12000 rpm at 4 °C for 10 min, and the supernatant was analyzed by LC-MS/MS (AB Sciex 4500 QQQ MS system connected with Shimadzu Nexera X2 LC system). All samples were qualitatively monitored for UNC9801 and aldehyde product. Data are reported in area ratio (ratio of UNC8732 and metabolites to Internal standard MS area response) at each time point for all samples.

### Generation of 293 Flp-In T-REx pools for BioID

293 Flp-In T-REx cells were co-transfected with pOG44 (Flp-recombinase expression vector) and a plasmid containing the coding sequence for FLAG-miniTurbo (FmT) fusion protein. Transfections were performed with Lipofectamine® 2000 (Invitrogen) according to the manufacturer’s instructions. After transfection, cells were selected with hygromycin at 200 ug/mL.

### BioID of NSD2 in 293 T-REx

Stable cell lines containing the NSD2-FmT were divided into two treatment conditions, with each condition containing 5x 15 cm plates at ∼80% confluency. Expression of the fusion protein was induced by 1 ug/mL tetracycline treatment overnight. The following day, the cells were treated with 10 µM MG-132 proteosome inhibitor (ab141003) and 5 µM UNC8732 or DMSO for 150 minutes, followed by the addition of biotin (50 µM final concentration) for 90 minutes. At the endpoint, the culture medium was removed, 2 mL PBS was added to each plate, and cells were harvested by scraping. Cells were collected in 15 cm tubes pelleted by centrifugation at 300×g for at least 5 minutes. The pellet was washed once with 10 mL of 1X PBS, then centrifuged at 300×g for at least 5 minutes to pellet the cells. PBS was removed, and the resulting cell pellet was flash-frozen in liquid nitrogen and immediately stored at -80 °C.

BioID^33^ was performed as described previously.^34^ In brief, cells were collected and pelleted (2,000 rpm, 3 minutes), the pellet was washed twice with PBS, and dried pellets were snap frozen. The cell pellet was resuspended in 10 ml lysis buffer (50 mM Tris-HCl pH 7.5, 150 mM NaCl, 1 mM EDTA, 1 mM EGTA, 1% Triton X-100, 0.1% SDS, 1:500 protease inhibitor cocktail (Sigma-Aldrich), 1:1,000 benzonase nuclease (Novagen)) and incubated on an end-over-end rotator at 4°C for 1 hour, briefly sonicated to disrupt any visible aggregates, then centrifuged at 45,000 x g for 30 minutes at 4°C. Supernatant was transferred to a fresh 15 mL conical tube. 30 µL of packed, pre-equilibrated streptavidin-sepharose beads (GE) were added and the mixture incubated for 3 hours at 4°C with end-over-end rotation. Beads were pelleted by centrifugation at 2,000 rpm for 2 min and transferred with 1 ml of lysis buffer to a fresh Eppendorf tube. Beads were washed once with 1 mL lysis buffer and twice with 1 mL 50 mM ammonium bicarbonate (ammbic, pH 8.3). Beads were transferred in ammbic to a fresh centrifuge tube and washed two more times with 1 mL ammbic. Tryptic digestion was performed by incubating the beads with 1 µg MS-grade TPCK trypsin (Promega, Madison, WI) dissolved in 200 µL of 50 mM ammbic (pH 8.3) overnight at 37°C. The following morning, 0.5 µg MS-grade TPCK trypsin was added, and beads were incubated an additional two hours at 37°C. Beads were pelleted by centrifugation at 2,000 x g for 2 min, and supernatant was transferred to a fresh Eppendorf tube. Beads were washed twice with 150 µL of 50 mM ammbic, and washes were pooled with the eluate. The sample was lyophilized and resuspended in buffer A (0.1% formic acid). 1/5th of the sample was analyzed per MS run.

#### Mass spectrometry

High performance liquid chromatography was conducted using a 2 cm pre-column (Acclaim PepMap 50 mm x 100 um inner diameter (ID)), and 50 cm analytical column (Acclaim PepMap, 500 mm x 75 um diameter; C18; 2 um; 100 Å, Thermo Fisher Scientific, Waltham, MA), running a 120 minutes reversed-phase buffer gradient at 225 nl/minute on a Proxeon EASY-nLC 1000 pump in-line with a Thermo Q-Exactive HF quadrupole-Orbitrap mass spectrometer. A parent ion scan was performed using a resolving power of 60,000, then up to the twenty most intense peaks were selected for MS/MS (minimum ion count of 1,000 for activation) using higher energy collision induced dissociation (HCD) fragmentation. Dynamic exclusion was activated such that MS/MS of the same m/z (within a range of 10 ppm; exclusion list size = 500) detected twice within 5 seconds were excluded from analysis for 15 seconds. For protein identification, Thermo .RAW files were converted to the .mzXML format using Proteowizard^35^, then searched using X!Tandem^36^ and COMET^37^ against the Human RefSeq Version 45 database (containing 36,113 entries). Data were analyzed using the trans-proteomic pipeline (TPP)^38^ via the ProHits software suite (v3.3).^39^ Search parameters specified a parent ion mass tolerance of 10 ppm, and an MS/MS fragment ion tolerance of 0.4 Da, with up to 2 missed cleavages allowed for trypsin. Variable modifications of +16@M and W, +32@M and W, +42@N-terminus, and +1@N and Q were allowed. Proteins identified with an iProphet cut-off of 0.9 (corresponding to ≤1% FDR) and at least two unique peptides were analyzed with SAINT Express v.3.6.^40^ Twenty control runs (from cells expressing the FlagBirA* epitope tag) were collapsed to the two highest spectral counts for each prey and compared to the two biological replicates (each with two technical replicates) of IRS1 BioID. High confidence interactors were defined as those with bayesian false discovery rate (BFDR) ≤0.01.

All mass spectrometry data are available at massive.ucsd.edu – accession: MSV000093206

### BioID Data Visualization

Proximity-dependent biotinylation data were analyzed in R (4.2.3) using limma (3.50.3). Data were filtered based on a Bayesian False Discovery Rate (BFDR) threshold of 0.01 to remove low confidence interactors. Spectral counts were normalized using the voom method. This process includes estimating the log-counts’ mean-variance relationship, generating precision weights, and performing a log2 transformation, thereby preparing the count data for linear modeling. A linear model was fitted to the normalized data using lmFit. The design matrix included UNC8732 treatment and DMSO as control. Technical replicate information was incorporated as a blocking factor in the model using duplicateCorrelation. This function estimates the correlation between technical replicates and uses it to adjust the standard errors of the log-fold changes, allowing separation of the technical variability (noise) from biological variability. Subsequently, empirical Bayes statistics were employed to moderate the standard errors of the estimated log-fold changes for each protein using the eBayes function. This enhances the reliability of the statistics by borrowing information across all proteins. Differential enrichment analysis was performed using makeContrasts with a contrast between UNC8732 treatment and DMSO control. P-values were adjusted for multiple testing using the Benjamini-Hochberg procedure. Proteins were deemed significantly differentially enriched based on a fold change threshold of 1.5 and an adjusted p-value threshold of 0.05. Sample clustering was done using the functions umap (umap library v0.2.7.0) and prcomp (stats library) in R on thirty-five significantly enriched proteins with an adjusted p-value < 0.05. Analysis code is available on GitHub at <https://github.com/d0minicO/NSD2_BioID>. The volcano plot was generated via the normalized spectral counts and adjusted p-value by GraphPad Prism 10.1.

### NanoBRET PPI Assay

NSD2 full-length template was cloned in frame into a pNLF1 vector (Promega) using the In-Fusion HD Cloning kit (Takara) to generate NanoLuc fusion on the N-terminus of NSD2. FBXO22 full-length template was similarly cloned into a pHTC-HaloTag vector (Promega) to generate HaloTag fusion on the C-terminus of FBXO22. The FBXO22-C326A mutant plasmids are generated via Q5 Site-Directed Mutagenesis Kit (NEB E0552) based on the wildtype FBXO22-HaloTag plasmid.

100 µL of resuspended U2OS cells at 4 × 10^5^ cells/mL were seeded per well in 96-well white opaque bottom plates. For each well, 0.01 µg of the NanoLuc-NSD2 plasmid, 0.03 µg of FBXO22-HaloTag plasmid, and 0.06 µg of empty vector pcDNA were transfected by XtremeGene HP (Millipore Sigma 6366236001) per manufacture instructions. After overnight transfection, the medium was completely removed, and 20 µL of 1:1000 (diluted in phenyl red-free DMEM with 4% FBS) HaloTag™ NanoBRET™ 618 Ligand (Promega G9801) was added to each well, followed by the addition of 20 µL of compound (or DMSO) at 2X concentration (diluted in phenyl red-free DMEM with 4% FBS). Upon mixing, the total volume is 40 µL per well with 1X compound. Two wells were reserved for no ligand control (40 µL of phenyl red-free DMEM only). The cells are incubated in a cell culture incubator for the prescribed time. At the endpoint, 10 µL of 1:100 (diluted in phenyl red-free DMEM with 4% FBS) Nano-Glo substrate (Promega N1572) was added in each well, and the plate was immediately scanned (with orbital mixing) on a CLARIOstar plate reader (BMGlabtech). The data collection and subsequent background subtraction follow the Promega technical manual (Literature # TM616). The background-subtracted BRET units were normalized to the control-treated wells (set = 1.00) from the same experiment.

### siRNA Knockdowns

Cells were seeded in 24-well format at 1 × 10^5^ cells per well. siRNAs targeting FBXO22 (Dharmacon siGENOME SMARTPool M-010812-01-0005), SKP1 (Dharmacon siGENOME SMARTPool M-003323-04-0005), CUL1 (Dharmacon siGENOME SMARTPool M-004086-01-0005) or Control siRNA (Millipore Sigma SIC001) were transfected using Lipofectamine^TM^ RNAiMAX Transfection Reagent (Invitrogen 13778075) per manufacture protocol. Cell culture medium containing transfection reagent was gently replaced the next day with fresh medium.

### Bio-Layer interferometry measurements

Bio-layer interferometry analysis was performed using an Octet RED384 instrument (ForteBio). C-terminally biotinylated SKP1-FBXO22 fusion proteins (WT or C326A) were immobilized onto SA biosensors (Sartorius) to a response level of 1 nm, then dipped into sample wells containing different concentrations of analyte. For analysis of compound-mediated ternary complex formation, NSD2-PWWP1 was held at a constant concentration of 2 μM while the concentration of compound (UNC10088 or UNC8732) was varied. All experiments were conducted at room temperature in commercial 1xHBS-EP+ buffer (GE Healthcare, 10 mM HEPES pH 7.4, 150 mM NaCl, 3 mM EDTA, 0.05% Surfactant P20) supplemented with 1 mM TCEP. Individual biosensors were used for each data point. Concentration-dependent steady-state binding analysis was performed using a one-site binding model in GraphPad Prism v9.0.1.

### Protein cloning, expression, and purification

#### Biotinylated NSD2-PWWP1 for SPR

Biotinylated NSD2-PWWP1 was expressed and purified as previously reported in Dilworth et al.^14^

#### His-NSD2-PWWP1

NSD2-PWWP1 (amino acids 211-350) was cloned into a pET28-MHL (RRID:Addgene_26096) vector with a N-terminal His-tag followed by a TEV-cleavage site. Plasmids were transformed into Escherichia coli BL21(DE3) CodonPlus cells and grown in Terrific Broth (Sigma) at 37°C. Once cells reached an OD600 ∼ 0.8, protein expression was induced by the addition of 0.5 mM IPTG at 16°C overnight. Cells were harvested by centrifugation (3000 x g) and purified as previously described.^14^

#### Sortaseable NSD2-PWWP1

Purified His-NSD2-PWWP1 protein was cut with protease TEV in buffer 20mM Tris, 500mM NaCl, 2mM TCEP, pH6.5 at the of ratio 1:20 (TEV: protein), 40°C overnight, followed by Talon beads incubation to remove TEV (His-tagged) and cut off His-tag. The resulting NSD2-PWWP1 protein has Glycine as the first amino acid at its n-terminus.

#### SKP1-FBXO22 fusion protein

The full-length SKP1-FBXO22 fusion protein with a 3xGGGS-linker was cloned into a pNic-Bio2 vector that encodes for an N-terminal His-tag as well as a C-terminal Avi-tag (GenBank: JF912191) using an In-Fusion HD Cloning System kit (Takara), following the manufacturer’s instructions. The SKP1-FBXO22 mutant (C326A) was generated using a Q5 Site-Directed Mutagenesis Kit (NEB, cat. E0552). Both SKP1-FBXO22 WT and C326A were expressed in BL21(DE3) CodonPlus cells and amplified in M9 minimal medium supplemented with 50 μM ZnSO4. For biotinylation, plasmids were co-expressed with a pCDF-BirA vector (GenBank: JF914075.1) in M9 minimal medium supplemented with 50 μM ZnSO4 and 40 μM biotin. Protein expression was induced overnight with 1 mM IPTG at 18°C. Cell pellets were resuspended in 20 mM Tris (pH 7.5), 500 mM NaCl, 2 mM TCEP, 5 mM beta-mercaptoethanol, 1 mM PMSF, 2 mM benzamidine, 5% glycerol, 0.1% Triton X-100, 5 mM Imidazole and 10 μM ZnSO4. The cells were lysed by sonication then purified using immobilized metal affinity chromatography (IMAC) resin charged with cobalt (TALON resin; Takara). Proteins were eluted with 200 mM Imidazole in buffer containing 20 mM Tris pH 7.5, 500 mM NaCl, 2 mM TCEP, 5 mM beta-mercaptoethanol, 1 mM PMSF, 2 mM benzamidine and 10 μM ZnSO4, then further purified by size-exclusion chromatography. Proteins were flash frozen in liquid nitrogen and stored at -80°C in a final storage buffer containing 20 mM Tris pH 7.5, 150 mM NaCl, 2 mM TCEP, 5 mM beta-mercaptoethanol, 1 mM PMSF, 2 mM benzamidine and 10 μM ZnSO4 until use. The degree of biotinylation was estimated using a streptavidin-band shift assay.

#### Co-expressed SKP1/FBXO22

Co-expressed SKP1 (full-length) and FBXO22 (aa.13-403, WT or C326A) were cloned into pFBOH-LICnotag and pFBOH-MHL vectors, respectively. All were amplified and expressed in Spodoptera frugiperda (Sf9) as previously described.^41^ Cell pellets were collected and stored in a buffer containing 50 mM Tris pH 7.5, 500 mM NaCl, 5% glycerol, 5 mM beta-mercaptoethanol, 1 mM TCEP, 1 mM PMSF, 0.5% NP40, 1:10,000 benzonase, and 5 mM benzamidine. Cells were lysed by several cycles of freeze-thaw and purified as described above.

### Differential Scanning Fluorometry

DSF measurements were taken using a Light Cycler 480 II instrument (Roche Applied Sciences). Protein samples were prepared in 20 µl containing 0.1 mg ml−1 SKP1/FBXO22, 0.9-100 µM compound, and 5x SYPRO Orange dye (Invitrogen, 5000x stock). Temperature scan curves were analyzed via dRFU/dT. Plots were prepared with the assistance of the online DSFworld application.^42^

### Hydrogen/Deuterium eXchange Mass Spectrometry (HDX-MS)

6 protein samples were prepared in buffer (20mM Tris (pH 7.5), 200mM NaCl, 5% glycerol, 1mM TCEP, 2mM Benzamidine): (i) 7.5 μM SKP1/FBXO22, (ii) 7.5 μM SKP1/FBXO22 + 7.5 μM PWWP1-NSD2, (iii) 7.5 μM SKP1/FBXO22 + 7.5 μM PWWP1-NSD2 + 75 μM UNC10088, (iv) 7.5 μM SKP1/FBXO22 + 75 μM UNC10088, (v) 7.5 μM PWWP1-NSD2, (vi) 7.5 μM PWWP1-NSD2 + 75 μM UNC10088. HDX labeling was performed in 87% D_2_O (10mM Phosphate Buffer pD 7.5, 150 mM NaCl) at 20 °C for 0, 1, 5 and 10 mins. Quenching took place for 2 mins using 100 mM Phosphate Buffer at a pH 2.5 at 0 °C. The quenched sample was loaded onto a mixed Nepenthensin-2:Pepsin (1:1) column (15 °C) and desalted on a short C18 column (0 °C) at 0.2 mL/min for 3 mins using a mobile phase A (0.1% formic acid in water). The trapped, desalted peptides were then reverse-phase separated using a long C18 column at 0.04 mL/min with a 12 min gradient from 35% to 85% of mobile phase B (0.1% formic acid in acetonitrile). The eluting peptides were electrosprayed and analyzed using the SELECT SERIES Cyclic IMS (Waters Corp., U.K.) in HDMSe mode. Leu-Enk was used for lockmass (m/z axis calibration). Peptides were identified using ProteinLynx Global Server 3.0.3. (Waters Corp., U.K.). HDX uptake was determined using DynamX 3.0 (Waters Corp., U.K.). Cumulative HDX differences (ΔHDX-MS) were calculated using Microsoft Excel and mapped onto the AlphaFold model of FBXO22 using PyMOL. For a cumulative ΔHDX-MS peptide signal to be considered statistically significant, it must have exceeded the cumulative error by >1%.

### In Vitro Ubiquitination Assay

#### Protein Expression and Purification for In Vitro Ubiquitination

For in-vitro substrate ubiquitination assays, CUL1-RBX1, Hhari, and CDC34B were prepared as previously described.^43^ CUL1-RBX1 neddylation was also performed as previously described.^44^

Uba1, UBCH5b, UBCH7, and Ub were purified from BL21 (DE3)-RIL cells using untagged constructs for UBCH5b and Ub and His-tagged construct for Uba1 and UBCH7. UBCH7-expressing cells were lysed into a 50 mM HEPES pH 8.0, 200 mM NaCl, 5 mM BME buffer. Lysate was purified by nickel affinity chromatography followed by cleavage of the His tag using a GST-tagged HRV3C protease. The cleavage reaction was stopped by passing over glutathione affinity chromatography to remove the protease, and the resulting cleaved protein was further purified via size exclusion chromatography. Uba1-expressing cells were lysed into a 50 mM Tris 7.6, 200 mM NaCl, 20 mM Imidazole, 5 mM BME buffer and purified via nickel affinity chromatography. His-Uba1 was dialyzed overnight to lower the salt concentration to 50 mM and subsequently purified by anion-exchange and size-exclusion chromatography. Cells containing untagged Ub were lysed into a 50 mM Tris pH 7.6, 200 mM NaCl buffer. Lysate was acid shocked by adding acetic acid to pH 4.5 and then spun down to remove precipitation followed by overnight dialysis into 25 mM NaOAc pH 4.5. The resulting protein was further purified by cation-exchange and size-exclusion chromatography. Cells expressing untagged UBCH5b were lysed into a 50 mM HEPES pH 7.0, 5 mM DTT buffer, and the resulting lysate was purified via cation-exchange and size-exclusion chromatography.

To create the fluorescent substrate, a sortaseable NSD2-PWWP1 construct was created. Sortase labeling was carried out overnight on ice by combining 40 µM substrate, 10 µM peptide, and 1 mM FAM-LPETGG fluorescent peptide. The sortase reaction was quenched by addition of 10 mM EDTA and buffer-exchanged 2x into a 20 mM Hepes pH 8.0, 500 mM NaCl, 1 mM DTT buffer using a Cytiva PD10 desalting column to remove excess fluorescent peptide. The fluorescently-labeled substrate was then passed over a Ni-NTA resin to remove sortase and analyzed via SDS-PAGE to confirm purity of the resulting product.

#### Substrate Ubiquitination Assays

Substrate ubiquitination assays were performed using the previously described fluorescent NSD2-PWWP1. The reactions were prepared by first creating a master mix of E1 (Uba1), E2 (UBCH5B or UBCH7), HHARI (only w/ UBCH7), CDC34B, Nedd8-conjugated CUL1-RBX1, and MgATP on ice in a BSA-containing buffer (20 mM HEPES pH 8.0, 200 mM NaCl, 0.5 mg/mL BSA). Individual reaction tubes containing the fluorescent substrate, Ubiquitin, and the indicated compound and SKP1-FBX022 to be tested were also prepared on ice in the same buffer. To start the reaction, a standard amount of master mix was added to each reaction, resulting in a final concentration of 0.1 μM E1, 0.5 μM of *NSD2-PWWP1 and compound, 1 μM of E2, HHARI, CDC34B, CUL1-RBX1, and SKP1-FBXO22, 100 μM ubiquitin, and 5 mM MgATP. The reactions were allowed to run at room temperature and were stopped after 15 minutes by quenching with SDS sample loading buffer. The quenched samples were separated by molecular weight by SDS-PAGE under non-reducing conditions using GenScript SurePAGE 4-12% gels. A GE Amersham Typhoon 5 was used to visualize fluorescent bands corresponding to ubiquitinated substrate on the resulting gel.

