## Extended Data Figures 1-4 for "Recruitment of FBXO22 for Targeted Degradation of NSD2"

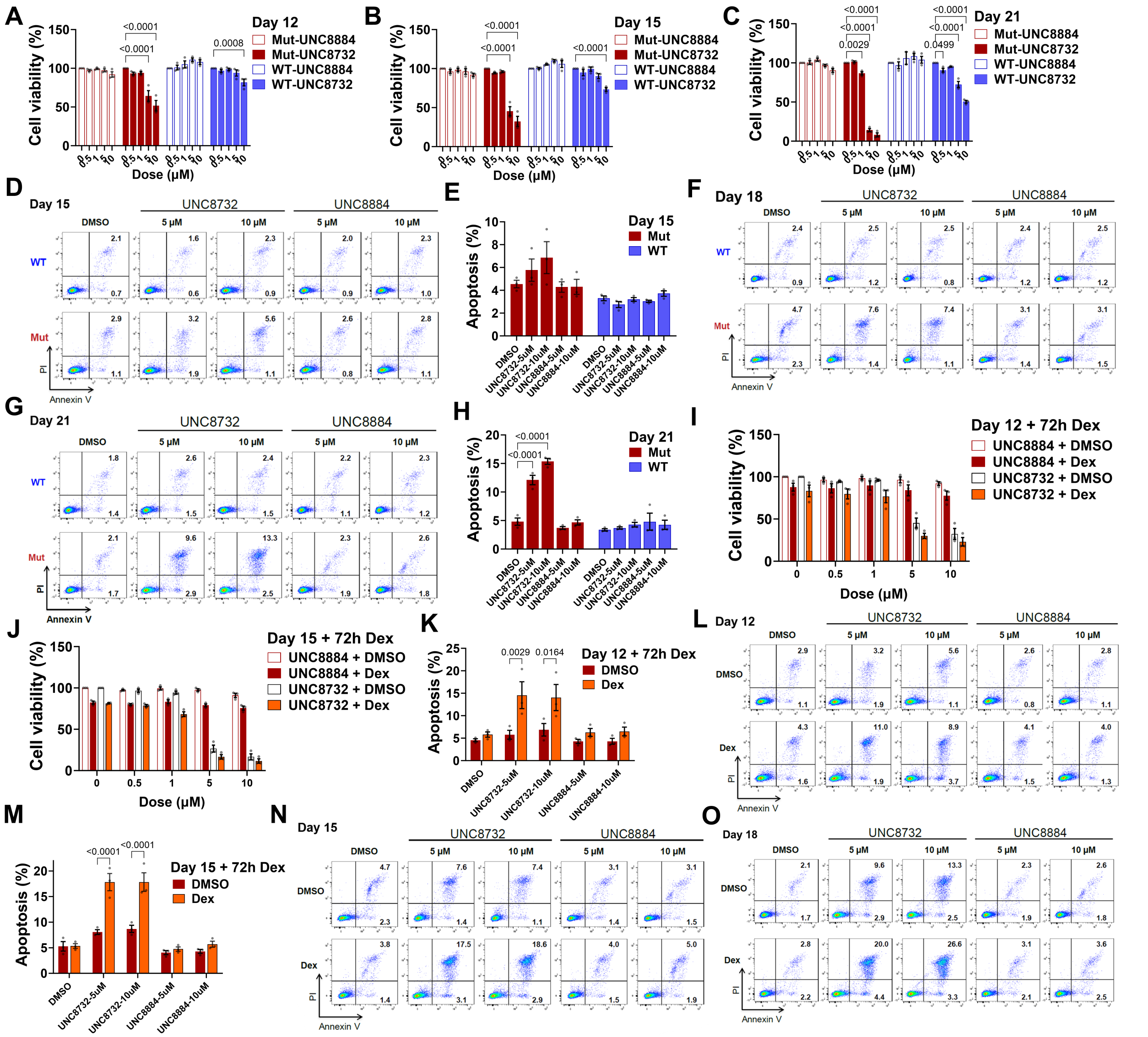


**Extended Data Figure 1**

**(A)** Viability of isogenic ALL cell lines RCH-ACV determined by CellTiter-Glo assay after treatment with varying concentrations of UNC8732 and UNC8884 for 12 days.

**(B)** Viability of isogenic ALL cell lines RCH-ACV determined by CellTiter-Glo assay after treatment with varying concentrations of UNC8732 and UNC8884 for 15 days.

**(C)** Viability of isogenic ALL cell lines RCH-ACV determined by CellTiter-Glo assay after treatment with varying concentrations of UNC8732 and UNC8884 for 21 days.

**(D)** Representative flow cytometric analysis plots of apoptosis of isogenic RCH-ACV cell lines treated with 5 μM and 10 μM of UNC8732 and UNC8884 for 15 days.

**(E)** Apoptosis of isogenic RCH-ACV cell lines detected using Annexin V/PI staining by flow cytometry after treatment with varying concentrations of UNC8732 and UNC8884 for 15 days.

**(F)** Representative flow cytometric analysis plots of apoptosis of isogenic RCH-ACV cell lines treated with 5 μM and 10 μM of UNC8732 and UNC8884 for 18 days.

**(G)** Representative flow cytometric analysis plots of apoptosis of isogenic RCH-ACV cell lines treated with 5 μM and 10 μM of UNC8732 and UNC8884 for 21 days.

**(H)** Apoptosis of isogenic RCH-ACV cell lines detected using Annexin V/PI staining by flow cytometry after treatment with varying concentrations of UNC8732 and UNC8884 for 21 days.

**(I)** Viability of *NSD2* mutant RCH-ACV cells determined by CellTiter-Glo after the pretreatment of varying concentrations of UNC8732 and UNC8884 for 12 days followed by dexamethasone (1 μM) for 72 hours.

**(J)** Viability of *NSD2* mutant RCH-ACV cells determined by CellTiter-Glo after the pretreatment of varying concentrations of UNC8732 and UNC8884 for 15 days followed by dexamethasone (1 μM) for 72 hours.

**(K)** Apoptosis of *NSD2* mutant RCH-ACV cell line detected using Annexin V/PI staining by flow cytometry after the pretreatment of varying concentrations of UNC8732 and UNC8884 for 12 days followed by dexamethasone (1 μM) for 72 hours.

**(L)** Representative flow cytometric analysis plots of apoptosis of *NSD2* mutant RCH-ACV cell line treated with 5 μM and 10 μM of UNC8732 and UNC8884 for 12 days followed by dexamethasone (1 μM) for 72 hours.

**(M)** Apoptosis of *NSD2* mutant RCH-ACV cell line detected using Annexin V/PI staining by flow cytometry after the pretreatment of varying concentrations of UNC8732 and UNC8884 for 15 days followed by dexamethasone (1 μM) for 72 hours.

**(N)** Representative flow cytometric analysis plots of apoptosis of *NSD2* mutant RCH-ACV cell line treated with 5 μM and 10 μM of UNC8732 and UNC8884 for 15 days followed by dexamethasone (1 μM) for 72 hours.

**(O)** Representative flow cytometric analysis plots of apoptosis of *NSD2* mutant RCH-ACV cell line treated with 5 μM and 10 μM of UNC8732 and UNC8884 for 18 days followed by dexamethasone (1 μM) for 72 hours.

Error bars represent mean ± SEM from three biological replicates. The statistical significance was evaluated using the Two-way ANOVA test and the p-values are displayed. WT, NSD2 WT; Mut, *NSD2* p.E1099K; Dex, Dexamethasone.

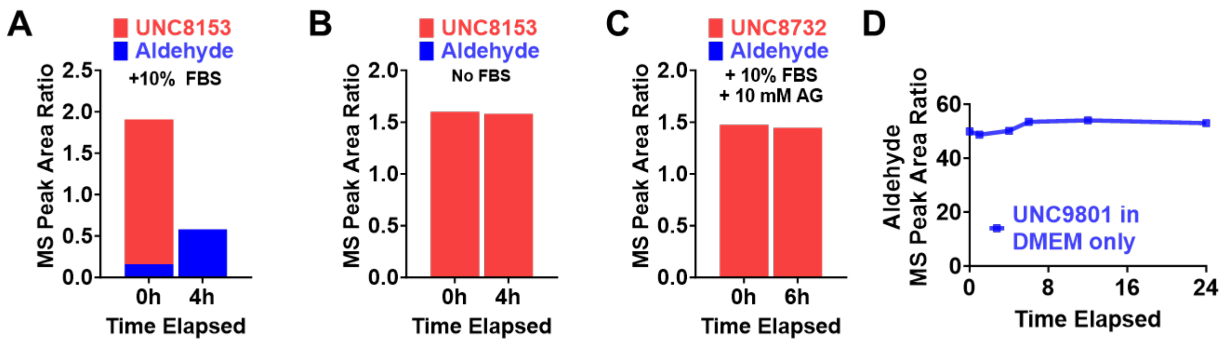


**Extended Data Figure 2**

**(A):** MS peak area ratio representing the relative levels of UNC8732 and its associated aldehyde species in cell-free DMEM + 10% FBS at indicated time points.

**(B):** MS peak area ratio representing the relative levels of UNC8732 and its associated aldehyde species in cell-free DMEM with no FBS at indicated time points.

**(C):** MS peak area ratio representing the relative levels of UNC8732 and its associated aldehyde species in cell-free DMEM + 10% FBS + 10 mM aminoguanidine (AG) at indicated time points.

**(D):** MS peak area ratio representing the relative level of the aldehyde adduct of UNC9801 upon addition of UNC9801 to DMEM + 10% FBS or DMEM only at indicated time points in hours.

* The experimental treatment was designed for 0h. However, due to the sample preparation process for MS, there's a brief period of simultaneous presence of the compounds in their respective conditions.

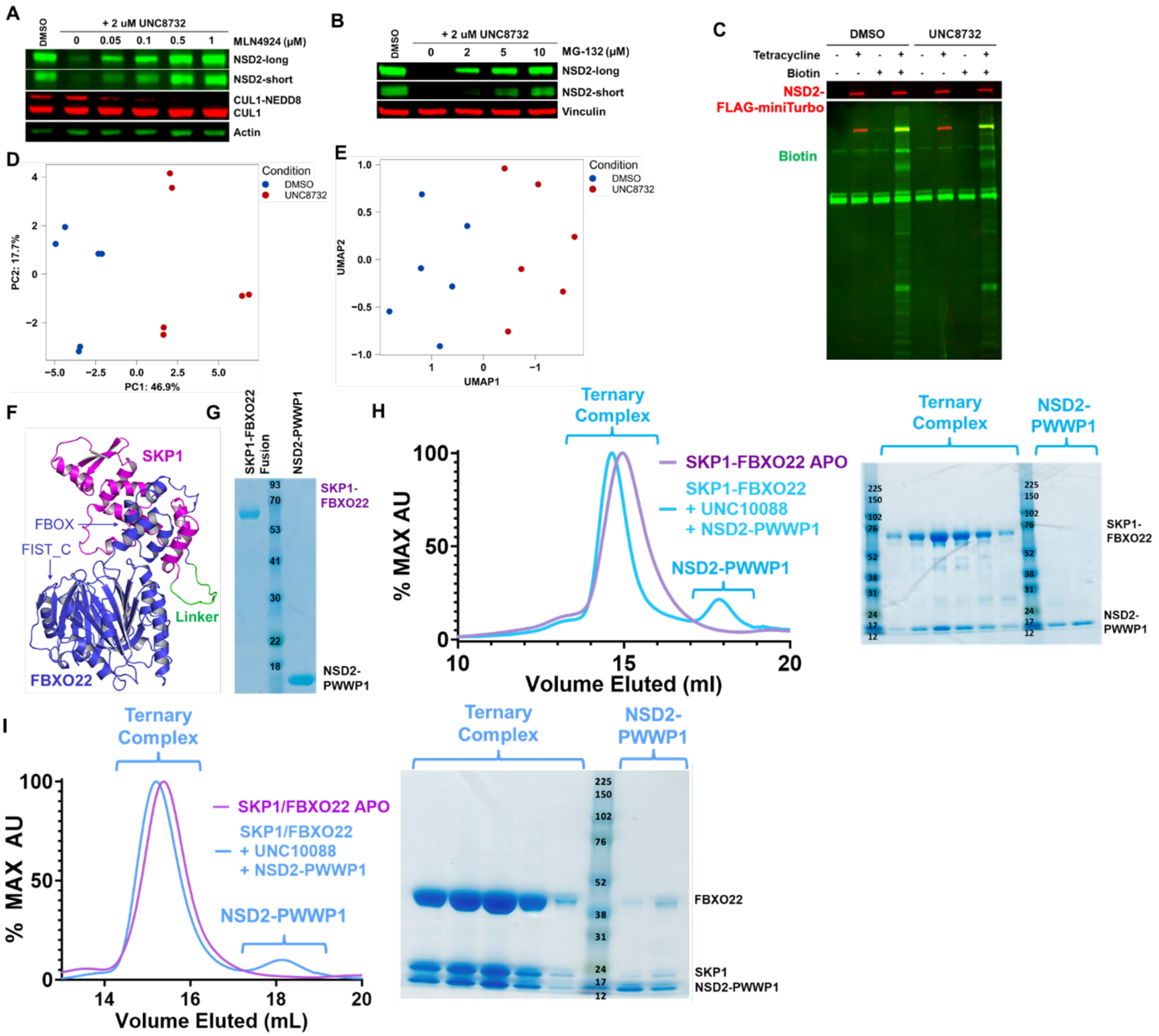


**Extended Data Figure 3**

**(A):** U2OS cells were co-treated with DMSO control or 2 μM of UNC8732 and indicated concentrations of MLN4924 neddylation inhibitor for 24h. Representative immunoblot shown. Experiment repeated independently three times with consistent results.

**(B):** U2OS cells were co-treated with DMSO control or 2 μM of UNC8732 and indicated concentrations of MG-132 proteosome inhibitor for 3h. Representative immunoblot shown. Experiment repeated independently three times with consistent results.

**(C):** T-Rex 293 cells stable cell lines with tetracycline inducible NSD2-FLAG-miniTurbo was treated with 10 μM MG-132 and the indicated combination of DMSO, 5 μM UNC8732, tetracycline (1 μg/mL) and biotin (50 μM).

**(D):** Principal Components analysis (PCA) of the BioID spectral counts data from DMSO (blue) or 5 μM UNC8732 (red) treated samples, each containing three independent experiments with 2 technical replicates for each independent experiment.

**(E):** Uniform Manifold Approximation and Projection (UMAP) analysis of the BioID data of DMSO (blue) or 5 μM UNC8732 (red), each containing 3 independent experiments with 2 technical replicates for each independent experiment.

**(F):** AlphaFold protein structure prediction of the SKP1-FBXO22 fusion protein. SKP1 is predicted to interact with the F-box domain of FBXO22 and the FIST_C domain remains somewhat separate for putative substrate interactions.
**(G):** SDS-PAGE analysis of the recombinant SKP1-FBXO22 fusion protein and the NSD2-PWWP1 post-purification.
**(H):** Left: Recombinant proteins of SKP1-FBXO22 fusion apo or SKP1-FBXO22 fusion + NSD2-PWWP1 + UNC10088 analyzed using a Superdex 200 Increase 10/300 GL column in the indicated combinations. Right: SDS-PAGE gel showing the eluted proteins of the SKP1-FBXO22 + UNC10088 + NSD2-PWWP1 condition at the indicated peaks.
**(I):** Left: Co-expressed SKP1/FBXO22 apo or co-expressed SKP1/FBXO22 + NSD2-PWWP1 + UNC10088 was analyzed using a Superdex 200 Increase 10/300 GL column in the indicated combinations. Right: SDS-PAGE gel showing the eluted proteins of the SKP1-FBXO22 + UNC10088 + NSD2-PWWP1 condition at the indicated peaks.

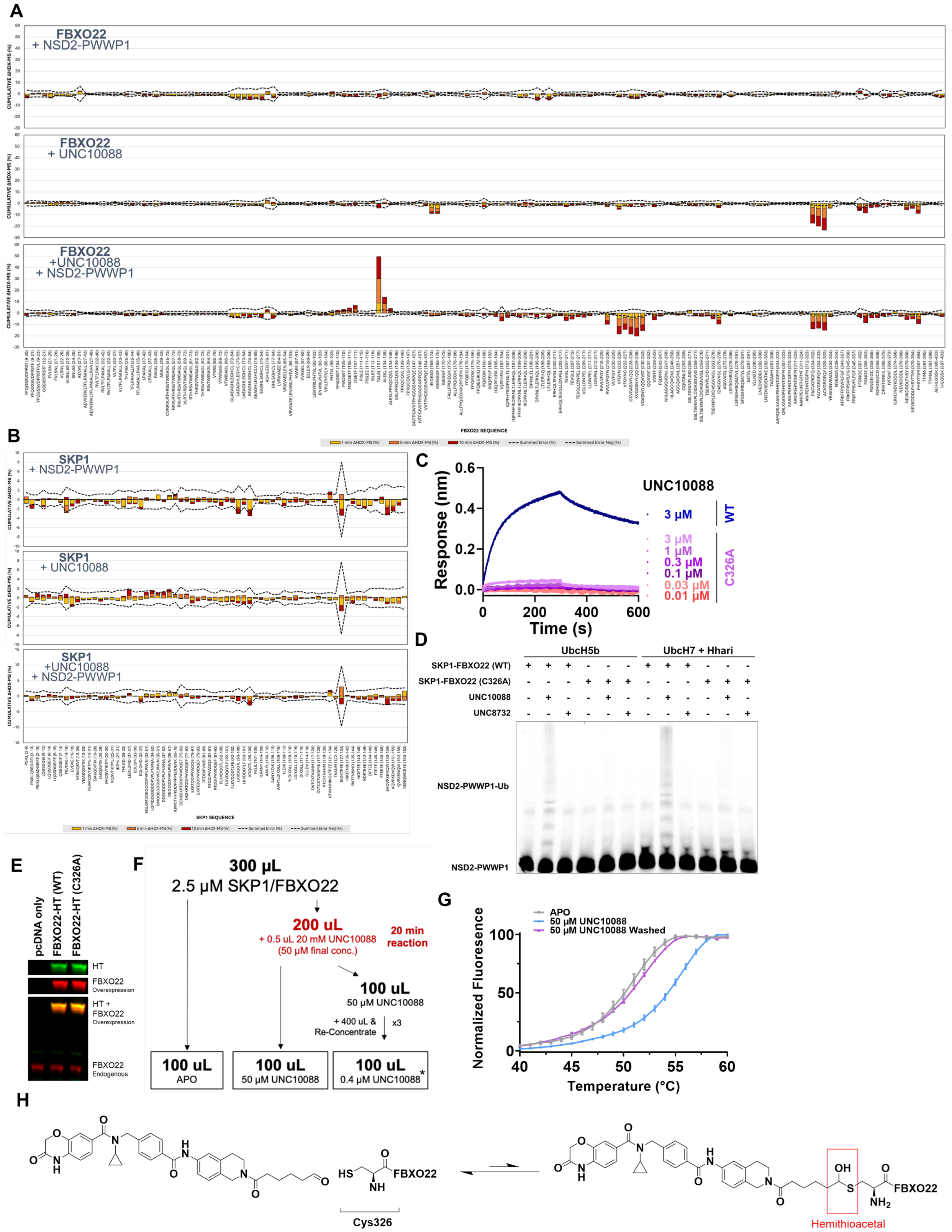


**Extended Data Figure 4**

**(A):** ΔHDX-MS of FBXO22. Co-purified SKP1/FBXO22 were incubated the presence of NSD2-PWWP1, UNC10088, or NSD2-PWWP1 + UNC10088. HDX was conducted at 1 (yellow), 5 (orange), and 10 minutes (red). To be statistically significant, the cumulative ΔHDX-MS signal must exceed the cumulative error (dashed line) by >1 %.

**(B):** ΔHDX-MS of SKP1. Co-purified SKP1/FBXO22 were incubated the presence of NSD2-PWWP1, UNC10088, or NSD2-PWWP1 + UNC10088. HDX was conducted at 1 (yellow), 5 (orange), and 10 minutes (red). To be statistically significant, the cumulative ΔHDX-MS signal must exceed the cumulative error (dashed line) by >1 %.

**(C):** Representative BLI sensorgrams upon the addition of increasing concentrations of UNC10088 and a fixed concentration of NSD2-PWWP1 (2 μM). WT or C326A-mutant SKP1-FBXO22 was loaded on SA biosensors to an average response of 1 nm. Curves are shown as the average of three independent experiments.

**(D):** In vitro NSD2-PWWP1 ubiquitination with the indicated sets of substrate priming machinery, UBCH5B or UBCH7 with HHARI. Each reaction also contained CUL1-RBX1 and CDC34B. The reaction mixture was analyzed by SDS-PAGE and fluorescence scanning. NSD2-PWWP1 was subject to a sortase reaction for fluorescent labeling. The gel presented is representative of three independent experiments.

**(E):** Immunoblotting following transfection of empty vector pcDNA, WT FBXO22-HT, or C326A-mutant FBXO22-HT in U2OS cells for 24h.

**(F):** Experimental flow chart for the designed 'wash' experiment to test for reversibility of the compound-FBXO22 interaction. * denotes estimated concentration based on the series of dilutions performed.

**(G):** Normalized fluorescence data from DSF for samples obtained from the experiment described in panel E. Error bars represent the SD of 3 technical replicates from one independent experiment.

**(H):** Schematic of the reaction between the aldehyde-containing degrader compound and the C326 of FBXO22.
